# Movement-Preceding Neural Activity under Parametrically Varying Levels of Time Pressure

**DOI:** 10.1101/2021.04.29.441753

**Authors:** Bianca Trovò, Yvonne Visser, Zafer İşcan, Aaron Schurger

**Affiliations:** Faculté des Sciences et Ingénierie, Sorbonne Université, Paris, France; Cognitive Neuroimaging Unit, INSERM U992, NeuroSpin, Frédéric Joliot Institute for Life Sciences, Commissariat à l’Énergie Atomique et aux Énergies Alternatives, Gif-sur-Yvette, France

## Abstract

Self-initiated movements are known to be preceded by the readiness potential or RP, a gradual increase in surface-negativity of cortical potentials that can begin up to 1 second or more before movement onset. The RP has been extensively studied for decades, and yet we still lack a clear understanding of its functional role. Attempts to model the RP as an accumulation-to-bound process suggest that this signal is a by-product of time-locking to crests in neural noise rather than the outcome of a pre-conscious decision to initiate a movement. One parameter of the model accounts for the imperative to move now, with cued movements having a strong imperative and purely spontaneous movements having no imperative. Two different variants of the model have been proposed, and both predict a decrease in the (negative) amplitude of the early RP as the imperative grows stronger. In order to test this empirically, we conducted an experiment where subjects produced self-initiated movements under varying levels of time pressure, and we investigated the amplitude, shape, and latency of the RP as a function of the imperative to move, operationalised as a time limit. We identified distinct changes in the amplitude of the early RP that grew non-linearly as the time limit grew shorter. Thus these data did not support the prediction made by the model. In addition, our results confirm that the shape of the RP is not stereotypically negative, being either positive or absent in about half of the subjects.

## Introduction

In everyday life the phenomenology of movement initiation can be reduced to two opposite case scenarios: movements are either exogenously induced in reaction to stimuli coming from the environment (for example a traffic light indicating the ‘go’ moment for crossing the intersection; a phone ring tone signaling when it is time to reply) or endogenously generated at an arbitrary undetermined time without any specific perceptual evidence (for example a football player kicking a penalty kick after waiting for the right moment; a computer user starting to browse for information on the internet).

A growing body of literature has been investigating this dichotomy between externally triggered and internally generated movements by contrasting cued responses originating from instructed acts with free, self-initiated ones [1–15] produced in the context of spontaneous voluntary movement (SVM) tasks [16]. Though useful in the context of laboratory experimentation, this polarization oversimplifies the real nature of movements which span a continuum from conditioned reaction time (RT) responses to waiting time (WT) responses (defined as the interval between the start of the trial and how long the subjects wait to freely initiate an act). Indeed, most real-life movements lie somewhere in between those extremes.

An attempt to provide a first mechanistic explanation of the neural phenomena lying along this spectrum was made by the stochastic decision model (SDM) [17], a leaky accumulator inspired by the class of integration-to-bound models in perceptual decision-making [18–22]. This model has managed to account for the distribution of waiting times and the shape of the readiness potential in a self-initiated movement task (a replication of Libet (1983) [23]). The readiness potential (RP) is a pre-movement buildup of neural activity traditionally identified as the electrophysiological sign of planning, preparation, and initiation of volitional acts [14, 15, 23].

The model interpreted the slowly building activity occurring before self-initiated movements both as a readout of the neural activity accumulating towards a decision bound and as a methodological artifact (‘selective biased sampling’) originating from averaging together data epochs aligned to the crests of auto-correlated neural noise [16, 17, 24, 25]. Furthermore, it posits that in the presence of a sensory signal from the environment (i.e., a strong imperative to act) it is mainly the neural activity triggered by the cue that determines the threshold crossing time. Conversely, lacking environmental information to accumulate, in the presence of only internal signals (i.e., a weak imperative to act) it is mainly the noisy background activity that determines the threshold crossing time.

Two variants of the model have thus far been proposed. One asserts a correspondence between the output of the accumulator process and the temporal profile of the RP [17]. The other asserts a correspondence between the movement-locked input to the accumulator and the temporal profile of the RP [26]. The output and the input model make opposite predictions regarding the direction of the relationship between the amplitude of the early RP and the waiting time but they both consider the imperative as a constant.

Here, we sought to manipulate the imperative reasoning that a progressive and continuous decrease in the signal triggering a movement would require a higher amount of background neural activity to accumulate in order to reach the threshold. To test this prediction we developed a new paradigm that allows us to study the initiation of movement along a spectrum between the extremes represented by ‘exogeneous’ and ‘endogeneous’ actions [1], laying the first stone on a new path of studies in volition focusing on continuous vs discrete events. This requires an innovative design for eliciting externally and self-initiated movements in continuous time.

So far, research on self-initiated movements has typically imposed a ‘minimal waiting time’ leading to a discrete movement initiation at a self-chosen time, such as the 3-25 s interval between irregular discrete self-paced movements in the Kornhuber & Deecke (1965) task [14] or the 3 s of rotating clock dial serving as buffer time before a discrete single self-initiated movement in the Libet (1983) task [23]. While a continuous Libet task, with parametrically varying minimal waiting times within which to perform the movement, has already been used for studying systematic effects on the RP [27], to our knowledge no design in the self-initiated paradigm has investigated movements made under a ‘maximal waiting time’ context, either in discrete or continuous time. The minimal waiting time design was developed with the goal of preventing rhythmic movements [14] or reaction type of movements [23] to occur in a freely initiated act. In our study we combined this minimal waiting time design (in the form of the traditional Libet clock-task) with a maximal waiting time one, by randomly increasing levels of temporal freedom (which is equivalent to decreasing levels of time pressure). Our goal was to indirectly manipulate the strength of the imperative to act by parametrically varying the time window in which participants were allowed to move. The external temporal constraint would be a proxy of the imperative to act by imposing a time pressure on the performance of a single self-initiated act.

We used EEG to record cortical readiness potentials from 22 healthy human participants while they performed the above-mentioned time limit task. As expected, self-chosen waiting times were longer with longer time windows (weaker imperatives to move), indicating that subjects were, on average, willing to wait longer when time permitted. EEG data revealed that the RP increased in amplitude for increasing levels of time pressure (i.e. shorter time windows), counter to the prediction of the model. These results were specific to the early phase of the RP, the one upon which the predictions of the SDM are hypothesized to apply: the late phase of the RP, the lateralized readiness potential (LRP) and the post-motor positivity (PMP) or reafferent potential, following the onset of the movement, did not yield any significant result.

While not consistent with the predictions of the model, these experimental findings show that the amplitude of the RP is susceptible to time pressure, and this puts some constraints on the viability of different theories of the origin of the RP. We discuss these findings in light of recent controversy surrounding the interpretation of the RP and the nature of the bounds in build-to-threshold decision models. We also suggest that future work focus on the spectrum from endogenous to exogenous determinants of movement initiation.

## Materials and Methods

### Participants

A total of 22 subjects participated in the experiment (3 females, median age 22 y, 1 left handed). All had normal or corrected-to-normal visual acuity. None had neurological or motor disorders. Subjects were recruited from the surrounding community and nearby universities through the CEA-GROOM online platform, and all subjects gave written informed consent to participate and were paid for their participation. The study was approved by the *Comité de Protection de Personnes Ile de France VII* in accordance with the Declaration of Helsinki (2008).

### Stimuli and Setup

Visual stimuli consisted of a circular clock dial (white on a black background) with a small fixation cross in the center (diameter= 5.5 cm, visual angle= 3.43°), a small dot revolving around the fixation cross along the inside edge of the clock dial, and a small tick mark indicating the time limit. The dot was red at the beginning of each trial, and would gradually change color to white over a period of 3 s. Subjects were instructed to perform the movement any time after the dot became completely white, but before the dot reached the tick mark. If subjects forgot to answer (missed trials) or the answer was too late or too early the screen would turn red and display the words: “Trop tard”, “Trop tôt”. Visual stimuli were back-projected onto a translucent viewing screen (Panasonic DLP projector, model PT-D7700E-K, 60 Hz refresh rate) positioned at ~100 cm in front of the subject’s eyes. Experimental stimulation was coded and presented with Psychtoolbox version 3 [28–30] running on MATLAB 2017 (R2017b, MathWorks Inc). Movements, consisting of finger lifts (index finger extension), were collected through a Fiber-Optic Response Pad (FORP; Science Plus Group) that sent a trigger pulse when the finger stopped breaking a beam of light. During the task the hand of the subject rested comfortably on a tabletop and on top of it the Fiber-Optic Pad was fixed with tape. The experimenter sat outside of the shielded room and communicated with the subject via an intercom. In order to ensure that participants followed the instructions correctly, at the end of the experiment we administered an anonymous questionnaire in which we asked subjects to report subjective impressions relative to their felt spontaneity during the task and possible sources of nuisance during the experiment (File S1).

### Design and Procedure

Each session began and ended with a 5-min resting-state recording (part of a separate study). During the actual experiment the subject had a training session lasting 10-min in which the experimental conditions were presented and re-explained. The two different types of conditions, resetting at each trial after 3 s of minimal waiting time, were: the time limit conditions, with constrained windows of time in which participants were allowed to make a movement (*2 s*, *4 s*, *8 s*, *16 s*) and the infinite condition (*Inf s*), an unlimited window of time like in the traditional Libet paradigm. After the instructions were clear, the recording would start with 10 blocks interleaved with 2-min breaks in between. During each block participants performed 20 finger lift movements, one per trial. Thus, there were a total of 200 trials in each experiment: max 40 randomized trials for each of the five conditions distributed across 20 trials in each of the 10 blocks. Depending on the subjects’ individual pace, the whole recorded session would last around one hour and no more than one hour and a half.

### Data Acquisition and Preprocessing

All data were acquired at NeuroSpin, in the French Alternative Energies and Atomic Energy Commission (CEA) center of Saclay, France. EEG recording was performed with the subject wearing an integrated 60-channel EEG cap (Elekta NeuroMag) at a sampling rate of 1,000 Hz (306-channel whole-head Elekta NeuroMag EEG/MEG system).The subject sat inside an electrically shielded chamber while they performed the tasks. An EEG reference was added on the tip of the nose and a ground electrode on the clavicle. The subject sat in an upright, but slightly reclined position. Electrooculogram (EOG) (horizontal and vertical) and electromyograms (EMG) (one above the flexor digitorum superficialis muscle and the other on the bone of the wrist, ulnar styloid process) were also recorded, using pairs of electrodes connected to bipolar recording channels. During the EEG cap preparation, we endeavored to keep impedance below 15 kΩ, while being mindful of any reported discomfort during the preparation. Data analysis was performed using MATLAB 2018 (MathWorks) with the help of the FieldTrip toolbox for MATLAB [31]. Dedicated trigger channels were used to insert temporal markers in the data, corresponding to: trial onset, button press, the five experimental conditions, fixation cross and clock appearance. For the average analysis, to preserve slow fluctuations, no detrending, no baseline correction, or high-pass filtering was performed. Data were down sampled to 250 Hz off-line prior to data analysis. Data were time locked to the finger lift response by epoching from 3 s before movement onset to 1 s after movement onset. Ocular and cardiac artifacts were isolated using Independent Components Analysis (ICA) [32] and removed by projecting the data onto the pseudoinverse of the artifact component matrix. Trials with significant artifacts remaining after this step were rejected based on visual inspection.

### Behavioral Analysis

After artifact rejection, we further cleaned trials from ‘cognitive artifacts’ [33] by removing trials in which participants committed errors (the behavioral response was ‘too early’ or ‘too late’ or the response was missing). This procedure was done after the EEG pre-processing in order to have a correspondence between the behavioral and electrophysiological trials. We tried to not exclude more than 20-25% of trials, thus keeping at least between 150-160 trials for each participant. The behavioral analysis served to ensure the success of the experimental manipulation: participants were expected to have on average increasingly slower waiting times with decreasing time pressure (or increasing ‘temporal freedom’). Therefore, to calculate descriptive statistics on the WTs we sorted the trials according to the time-limit condition.

### EEG Analysis

After rejecting trials with behavioral artifacts, we applied a fourth-order Butterworth low pass filter (30 Hz cutoff). Since time locking to high-pass–filtered EMG signal simply shifts the signal by ~50 ms forward in time as compared to time locking to the button press (or in our case finger lift) [17, 34], we chose to time lock to the finger lift by epoching from 3 s before movement onset to 1 s after movement onset. We performed data pre-processing in four different ways in order to adapt to the needs of the data analysis and visualization: with all the trials averaged together and all the conditions randomized; keeping individual trials and all the conditions randomized; by grouping the data into the five experimental conditions (*2 s*, *4 s*, *8 s*, *16 s*, *Inf s*) with all the trials averaged together; by grouping the data into the five experimental conditions and keeping each trial separate (for further details see code available in the GitHub repository). For the averaging analysis we mostly relied on the data grouped into the five experimental conditions. For the statistical analysis (regression) we used the pre-processed data with non averaged individual trials.

An important criterion for participant inclusion in the statistical analysis was the presence of a clear readiness potential. The RP is localized within the midline centro-parietal area [35]. Traditionally, the RP exhibits maximal amplitude in the electrode at the vertex (Cz or FCz) and this is the commonly used channel, along with lateral central electrodes contralateral to the hand used to perform movements (FC1, C1, C3) [34, 36]. However, the inter-subject variability of the RP amplitude at the candidate and neighboring electrodes sites makes it problematic to select a single electrode as the channel for the analyses. Indeed, most subjects exhibit an RP at electrode Cz and one or more adjacent electrodes, especially contralateral to the dominant hand used to perform the task, but the center of the spatial distribution varies from subject to subject (see [17], p. E2911). Visual inspection of the averaged RP’s time series at candidate electrodes (single sensor selection method) for each individual subject is one of the traditional ways to establish the presence of an RP but not necessarily the most optimal [37]. In the literature, indeed, there is no agreed-upon standardized procedure for assessment of the RP [38].

In order to obtain a more robust estimate of the RP, we customized a criterion-metric inspired by the effect-matched spatial filtering (EMSF or SF) method [39]: we computed the mean signal amplitude within a 50 ms window positioned at the pre-movement peak of the signal, and the mean signal amplitude within a 50 ms window placed 1 s earlier. We computed the difference between these two means and an amplitude decrease of at least 1*μV* was the criterion for considering the movement preceding activity to be a RP. Participants who showed a decrease that was inferior to this criterion or instead an increase in voltage prior to movement were excluded from EEG signal analysis.

Finally, to compute the Lateralized Readiness Potential we computed the average of the sensors C3 and C4 and then substracted their amplitude [40].

### Data Transformation and Statistics

To analyze differences in neural amplitudes we employed Wilcoxon’s signed rank test and linear regression (MATLAB function *regress*). To overcome the problem of applying a linear regression on autocorrelated EEG time points, the function regress was run for each time point independently, across all 22 subjects, across all 60 channels (see [41] chap. 34), across 2001 time points (for the latency −3 s 1 s), across all trials. The data dimension was: ~177 trials × 60 channels × 2001 time points. The function returned a vector of beta weights or regression coefficients (slopes and intercepts) whose dimension was 2 × 60 × 2001. For the predictor data, we specified a variables-by-observation design matrix X, with rows of X corresponding to observations, and columns corresponding to predictor variables. For the behavioral data the predictor (x) was the time limit condition indexes and the criterion variable (y) was the participant’s WTs on each trial. Given the non-linear relationship between the behavioral responses (subjects’ waiting times) and the time limit conditions, to allow for a linear regression, we log transformed the input variables.

In the time-series domain, we ran a trial-by-trial regression with each of the time limit conditions (*2 s*, *4 s*, *8 s*, *16 s*, *Inf s*) as a predictor (x) and the RP amplitudes at each time point as response variables (y) (henceforth *first regression*) across each of the subjects (n= 22). Statistical significance of the resulting 22 beta coefficients was then tested against zero via non-parametric approach with Fieldtrip’s cluster-based permutation test (function: *ft timelockstatistics*) that corrects for multiple comparisons without making any prior assumptions regarding where the effect should be. However, this function, for the paired t-test, allows only for testing two independent conditions against each other. Therefore, in order to test our regression coefficient against zero we created a template of 22 surrogate beta weights with value 0. To calculate the significance probability, we used the Monte Carlo Method [42] with dependent sample t-statistics. The cluster-based permutation tests were performed on the time intervals 2 s to −0.2 s relative to the finger lift and −0.2 s to 1 s after the finger lift with 1000 iterations from the permutation distribution. The cluster alpha value was set to 0.05 and we ran the cluster based permutation test under a two-tail hypothesis. Channel neighbours for spatial clustering were defined using the distance method (minimum neighbourhood distance = .13). We also ran a trial-by-trial regression with the behavioral WTs as predictors (x) and the RP amplitudes as response variables (y) (henceforth *second regression*). This second regression was performed as a control to test for the relationship between the WTs and the RPs. To correct for multiple comparisons, we followed the same procedure as outlined above.

### Computational Model

The computational model that we used was the same as that used in Schurger et al. (2012) [17] and Schurger (2018) [26], which amounts to numerical integration over the following differential equation:

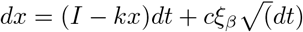

where x is the decision variable, I is the imperative or evidence, k is leak (exponential decay in x), *ξ_β_* is noise with 1*/f* exponent beta, and c is a noise scaling factor set at 0.1. For *β* = 0, *ξ_β_* is simply Gaussian white noise, and the process is a leaky stochastic accumulator (Ornstein-Uhlenbeck process). For *β* > 0, *ξ_β_* is autocorrelated (pink) noise. There is one additional parameter which is a threshold on x. When the threshold is crossed we consider that a movement has been initiated and we time lock the simulated data to that point in time. Two assumptions about the RP are possible. One is that the RP reflects the average trajectory of the decision variable leading up to threshold crossing [39], and the other is that the RP reflects the average time course of the noise (*ξ_β_*) leading up to the threshold crossing [26]. We consider both possible interpretations here, in case the predictions that they made were different (though it turns out that they were qualitatively the same).

The goal of the simulations was to make a prediction about the relative amplitude of the RP under different levels of imperative. To that end, we ran the simulation many times for a range of different values of I (the evidence or imperative) and observed how the amplitude of the simulated RP changed with changing I. We did this for both the input and output assumptions and also across a broad range of values for the other parameters, except for beta which was either 0 (output assumption) or 1.4 (input assumption). Repeating the simulation for a range of different values of k and for different thresholds reassured us that the prediction about the relative amplitude of the RP for different values of I was not limited to a very narrow range of values for the other parameters, and it turns out that it was not. Regardless of the specific value of k and the threshold (within reason - for extreme values the model simply breaks down) the same qualitative prediction emerged: the amplitude of the (early) RP is predicted to be higher for weak imperative and vice-versa. This was true under both the input and output assumptions. Figure 2 shows the results of the simulation for representative parameter values (see the figure legend for the specific parameter values).

**Fig 1.**
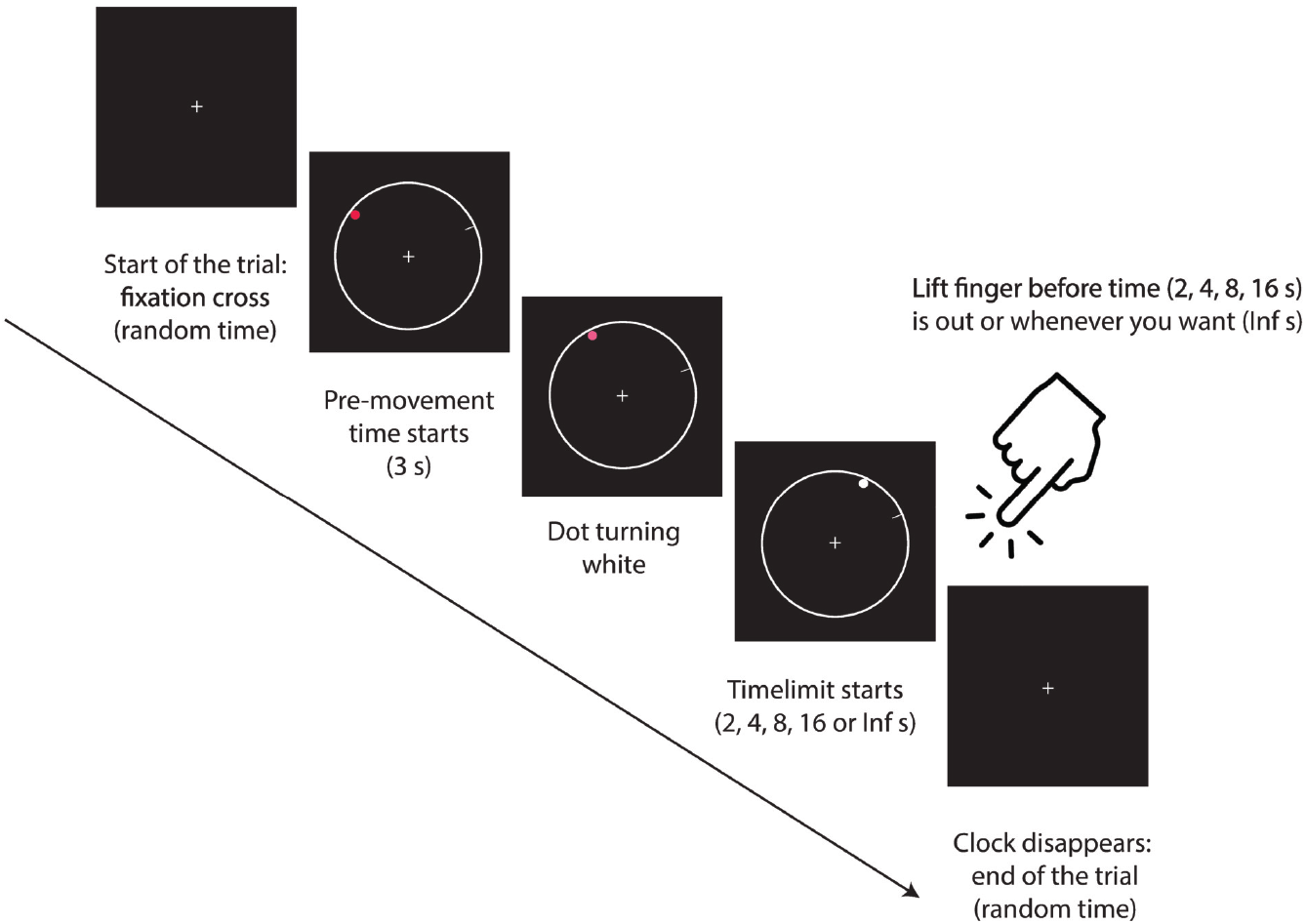
Timeline for a single trial. Participant watched a fixation cross while a dot revolving around the edge of the clock dial appeared. Once this dot completely turned white, the participant was allowed to make a finger lift any time between then and when the dot reached the tick mark (2, 4, 8 or 16 seconds later) or whenever they wanted if there was no tick mark (Infinite seconds). After a random delay following the finger lift, the trial ended).

**Fig 2.**
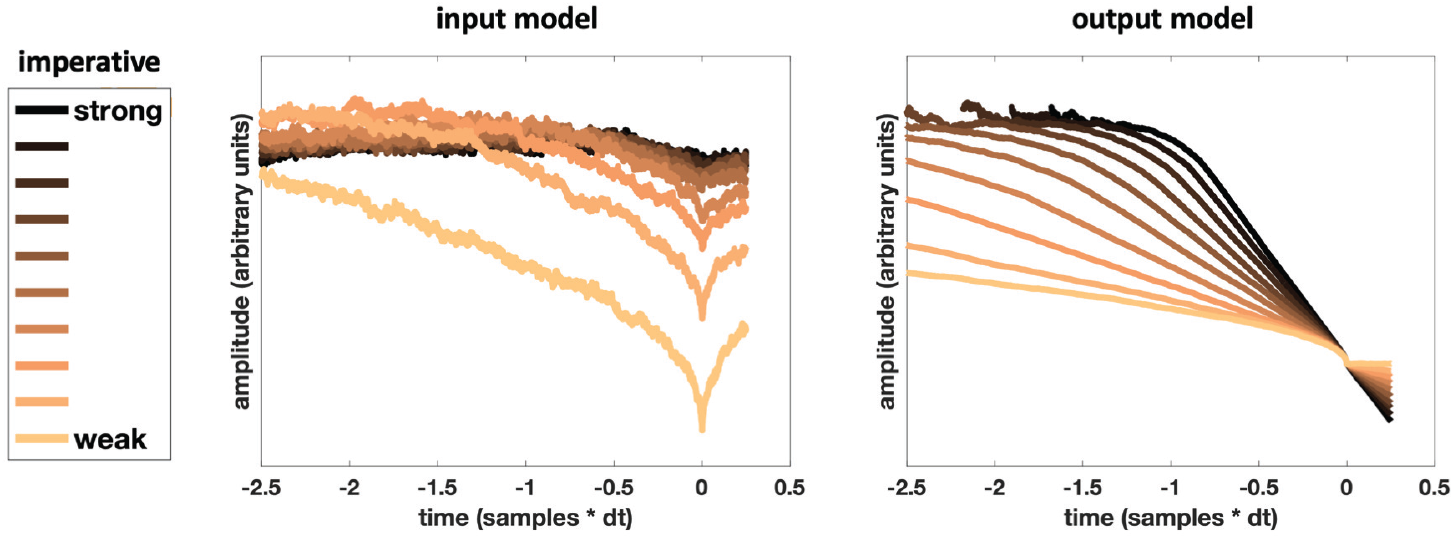
Model-driven predictions. **(A)** Average stochastic input to the accumulator, time aligned to first crossing times in the output, separately for a range of different values of I (the evidence or imperative to move) sorted by levels of strength. Darker lines (from brown to black) correspond to the shortest time limit conditions. Lighter lines (from yellow to orange) correspond to the longest time limit conditions. **(B)** Same as **(A)** but for the output of the accumulator.

## Results

### Behavioral Results

Participants (n = 22) were presented with a rotating dot on a clock dial while maintaining fixation in the middle of the clock. On each trial one of five different temporal windows of opportunity to make a movement (‘time limits’) was marked on the clock dial in the form of a small tick mark marking the end of the interval. We used a randomized design in which at each trial a different window of time was assigned from among the following: *2 s*, *4 s*, *8 s*, *16 s*, *Inf s* (see Methods). In principle, with the infinite time limit condition subjects could wait forever to perform a movement. In practice, due to the demand characteristics of the task, participants never waited more than 30 s to make a finger lift, like reported by Schurger et al. [17]. We received questionnaire answers for 20 participants out of 21. Only five of them reported adopting a particular strategy for performing the task. Six of them reported that the time limit triggered them to move, specially in the case of short duration. Four of them reported that the cue turning white triggered them to move faster or to wait before feeling the need to move. Fourteen of them reported that their actions were spontaneous most of the time, four that their actions were always spontaneous and only one of them said that their actions were only sometimes spontaneous.

Overall participants completed on average 185 (SD= 10.70) trials in the entire session, 177 (SD= 10.96) for correct trials only. The waiting-time data were log-transformed because they were markedly positively skewed (one-sample Kolmogorov-Smirnov test for normality: 0.787, *p* < 1.0*e* − 07; see Fig. S 1) and the standard deviation scaled with the mean (r= .88, *p* < 0.00001; see Fig. S 2), as is the case for response-time tasks [43]. Participants’ median waiting times within each condition, for correct trials only, were: 0.48 s (min: 0.002 s, max: 1.76 s; IQR: 0.39 s - 0.57 s) for time limit *2 s*; 1.31 s (min: 0.002 s, max: 3.7 s; IQR: 1.10 s - 1.53 s) for time limit *4 s*; 2.34 s (min: 0.05 s, max: 7.88 s; IQR: 1.77 s - 3.03 s) for time limit *8 s*; 3.60 s (min: 0.05 s, max: 15.11 s; IQR: 2.21 s - 4.98 s) for time limit *16 s*; 4.10 s (min: 0.01 s, max: 33.41 s; IQR: 1.70 s - 5.57 s) for the *Inf* condition. The Kruskal-Wallis Test, a non-parametric one-way ANOVA, revealed a significant main effect of the time limit durations assigned (*2 s*, *4 s*, *8 s*, *16 s*, *Inf*) on the median waiting times (H = 71.98, *p* < 9*e* 15, df = 4); post-hoc pairwise comparisons from a multiple comparison test (Mathworks; function *multcompare*) revealed that participants waited significantly less time in the *2 s* condition as compared to all other conditions, and waited significantly less time in the *4 s* condition compared to the conditions *16 s* and *Inf s* (for H statistics and p-values see Table 1 in Supporting Information). Participants’ average waiting times were highly correlated with increasing time limit conditions (R= 0.60, *p* < 0.0001; see 3a). Taken together these results show that participants waited longer when they were given more time, as expected.

**Table 1.** Criterion for deciding if a participant shows a canonical readiness potential.

| Subject n° | RP amplitude ($\mu V$) | RP peak latency (s) | SF baseline window (s) | Inclusion |
| --- | --- | --- | --- | --- |
| 1 | -2.775e-07 | 0.014 | [-0.036 0.014] | No |
| 2 | -2.872e-07 | -0.020 | [-0.070 -0.020] | No |
| 3 | -3.424e-06 | 0.128 | [0.078 0.128] | Yes |
| 4 | -4.306e-07 | 0.038 | [-0.012 0.038] | No |
| 5 | -9.340e-07 | -0.042 | [-0.092 -0.042] | No |
| 6 | -3.5671e-06 | 0.018 | [-0.032 0.018] | Yes |
| 7 | -2.400e-06 | -0.030 | [-0.080 -0.0300] | Yes |
| 8 | -1.593e-06 | -0.190 | [-0.240 -0.190] | Yes |
| 9 | -2.605e-07 | -0.200 | [-0.250 -0.200] | No |
| 10 | -1.607e-06 | -0.140 | [-0.190 -0.140] | Yes |
| 11 | -6.058e-07 | 0.056 | [0.0060 0.0560] | No |
| 12 | 1.029e-06 | -0.200 | [-0.250 -0.200] | No |
| 13 | -2.702e-06 | -0.016 | [-0.066 -0.0160] | Yes |
| 14 | 1.070e-06 | 0.022 | [-0.028 0.022] | No |
| 15 | -2.711e-06 | -0.096 | [-0.146 -0.096] | Yes |
| 16 | 1.775e-06 | -0.0880 | [-0.138 -0.088] | No |
| 17 | -2.344e-06 | -0.002 | [-0.052 -0.002] | Yes |
| 18 | -1.486e-06 | -0.050 | [-0.100 -0.050] | Yes |
| 19 | -1.596e-06 | -0.200 | [-0.250 -0.200] | Yes |
| 20 | -3.074e-06 | -0.054 | [-0.104 -0.054] | Yes |
| 21 | -3.856e-06 | 0.110 | [0.060 0.110] | Yes |
| 22 | 1.463e-07 | -0.172 | [-0.222 -0.172] | No |

Criterion defined as a decrease of minimum $1\mu V$ over 1 s ( $-1e-06$ ). RP stands for Readiness Potential, SF stands for Spatial Filter.

Given the temporal aspect of the task, as a post-hoc analysis we wondered if participants would adopt a different strategy in response to the time limit conditions. Because the distribution of median WTs of each subject across condition did not show any segregated pattern, we decided to look at their standard deviation averaged across conditions (see Fig. S1, A and B). Following what the literature has described as a ‘universal law’ of reaction time tasks [20, 43] or timescale invariance [44, 45], we showed that most of the participants would display more random responses for longer time limits, and that their WTs would become more dispersed with increasing temporal freedom or decreasing temporal pressure (Fig. S1, C). By looking at within-subjects standard deviation responses, we found 15 participants (subjects 2, 3, 6, 7, 9, 11, 12, 13, 14, 15, 17, 18, 19, 20, 21) whose standard deviation linearly scaled with time limit imposed. We called those ‘timers’. The other 7 participants (subject 1, 4, 5, 8, 10, 16, 22), instead, displayed very little change in variability across conditions (meaning that the standard deviations of their WTs did not scale with their mean WTs). We considered those subjects as Non-timers because they were not sensitive to the timing aspects of the task, as opposed to the Timers. We wondered if this difference in the behavior would reflect some neural process related to the RP and potentially connect the RP to an endogenous timing mechanism. Therefore, we later used this categorization across participants for sorting the electrophysiological data into two distinct groups for group-level statistical analysis and for comparison with a subset of participants displaying different temporal profiles of RP (see next session).

### The Readiness Potential is Not a Stereotypical Signal Across Subjects

Before undertaking any group-level analysis on the electrophysiological data we wanted to make sure that all the included participants displayed a clear RP, which is canonically defined as a negative-going voltage deflection before movement onset with a fronto-central scalp distribution. We then performed a within-subject analysis: we examined the EEG time-series by time locking the data to the onset of movement (finger lift), sorted the trials according to each of the five experimental conditions and took the average of the samples over 4 s (epoch: −3 s to 1 s). We looked at those individual mean amplitudes with a customized criterion for determining the presence of the RP across conditions (see Methods for details) using as a reference the infinite time limit condition because its RP is obtained through the classical Libet task. Through this technique participants were divided into two groups: the ones showing a canonical negative-going RP signal and the ones showing a positive-going RP signal or a flat RP signal (see Fig. S2). The last two groups represented up to 45.5% of the entire sample size. With the criterion method we concluded that only 12 subjects (subjects 3, 6, 7, 8, 10, 13, 15, 17, 18, 19, 20, 21) exhibited a stereotypical RP in the *Inf s* condition. In 10 subjects (subjects 1, 2, 4, 5, 9, 11, 12, 14, 16, 22) the RP could not be identified at all (see Table 1). We grouped these participants in the following categories: Negative-RPs and Positive-RPs. We later used this categorization across participants for post-hoc analysis. Interestingly, we found that the morphologies of the ERPs of the RP-negative subjects and the ones of the timers overlapped, as well as the ERPs of the RP-positive subjects and the ones of the non-timers (see Fig. S3 a) and b), c) and d)).

We computed the grand average of the time-locked data across all the 22 participants around three RP candidate sensors (Cz, FCz and C3)(see Fig. 4a). Plotting the between-subjects mean RP amplitudes as a function of condition monotonically ordered (*2s*, *4s*, *8s*, *16s*, *Inf*) revealed that the mean RP amplitudes across participants decreased with increasing time limits, meaning that amplitude was negatively correlated with the imperative (see Fig. 3b), contrary to the prediction of the model (see Fig. 2). We initially looked for differences in mean amplitude between conditions with Wilcoxon signed-rank tests and corrected for multiple comparisons. We found no significant difference between adjoining conditions, and only the *2 s* time limit condition significantly differed from the others (see Table S 3). Given the clear non-linear monotonic trend in the electrophysiological data, we decided to perform a regression analysis in order to see if the RP signal decreased or increased as a function of the imposed time limit condition (henceforth *first regression*). Furthermore, as a control we also tested if the RP signal decreased or increased as a function of the WT (henceforth *second regression*).

**Fig 3.**
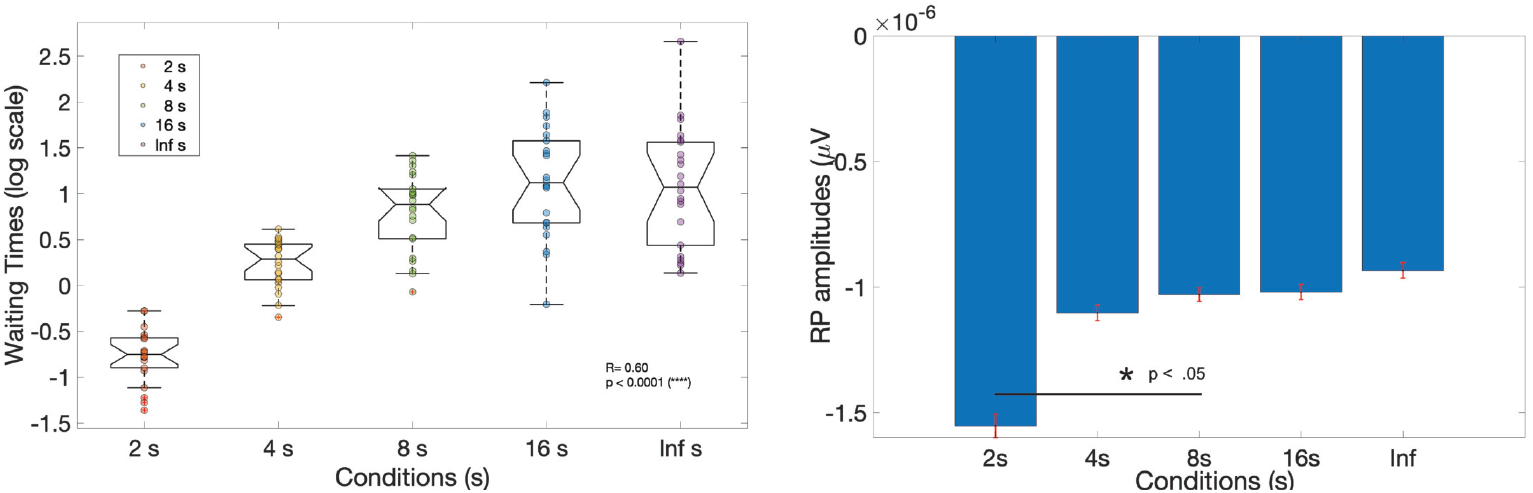
Behavioral performance and RPs sorted by condition. **(A)** Average waiting times sorted by conditions 2*s* (red dots), 4*s* (yellow dots), 8*s* (green dots), 16*s* (blue dots), *Inf* (violet dots) and plotted in log scale. Dots represent individual participants (n= 22). Mid line in box plots indicate the median WTs across subjects. **(B)** Average RP amplitudes across participants (in blue) sorted by conditions 2*s*, 4*s*, 8*s*, 16*s*, *Inf*; small red bars represent ± SEM across participants.

**Fig 4.**
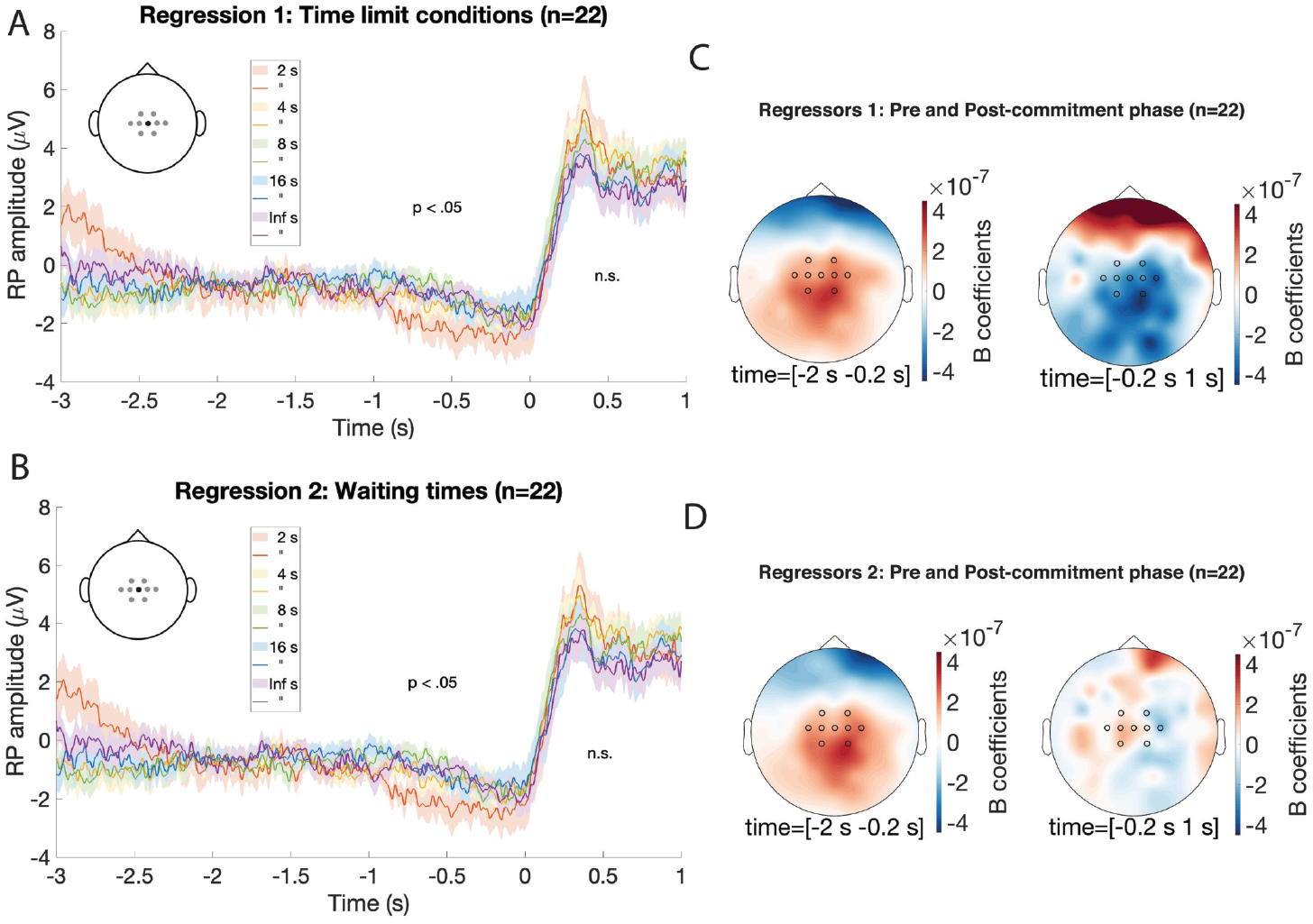
**(A, B)** Grand-averaged RP amplitude SEM across participants for the conditions 2*s* (red line and shade), 4*s* (yellow line and shade), 8*s* (green line and shade), 16*s* (blue line and shade), *Inf* (violet line and shade). Data are time-locked to time 0 s (finger lift). For representative purposes, we display EEG activity recorded from Cz electrode which did not differ from the ROI cluster of channels. P-values indicates cluster analysis period for the main hypothesis. In **(A)** cluster-based permutation testing was used to test whether the regressions of single-trial RP amplitudes against the time limit conditions per each trial (predictors in regression 1) was significant. In **(B)** cluster-based permutation testing was used to test whether the regressions of single-trial RP amplitudes against the single-trial WT responses (predictors in regression 2) was significant. **(C, D)** Topography of the grand-averaged beta coefficients’ power for regressions using the time limit conditions per each trial as predictors **(A)** or the single-trial WT responses as predictors **(B)**. The time interval (s) is indicated below each subplot and corresponds to the the early RP or pre-commitment phase [−2 s to ~−0.2 s] and the late RP [~−0.2 s to 0 s] or post-commitment phase.

**Fig 5.**
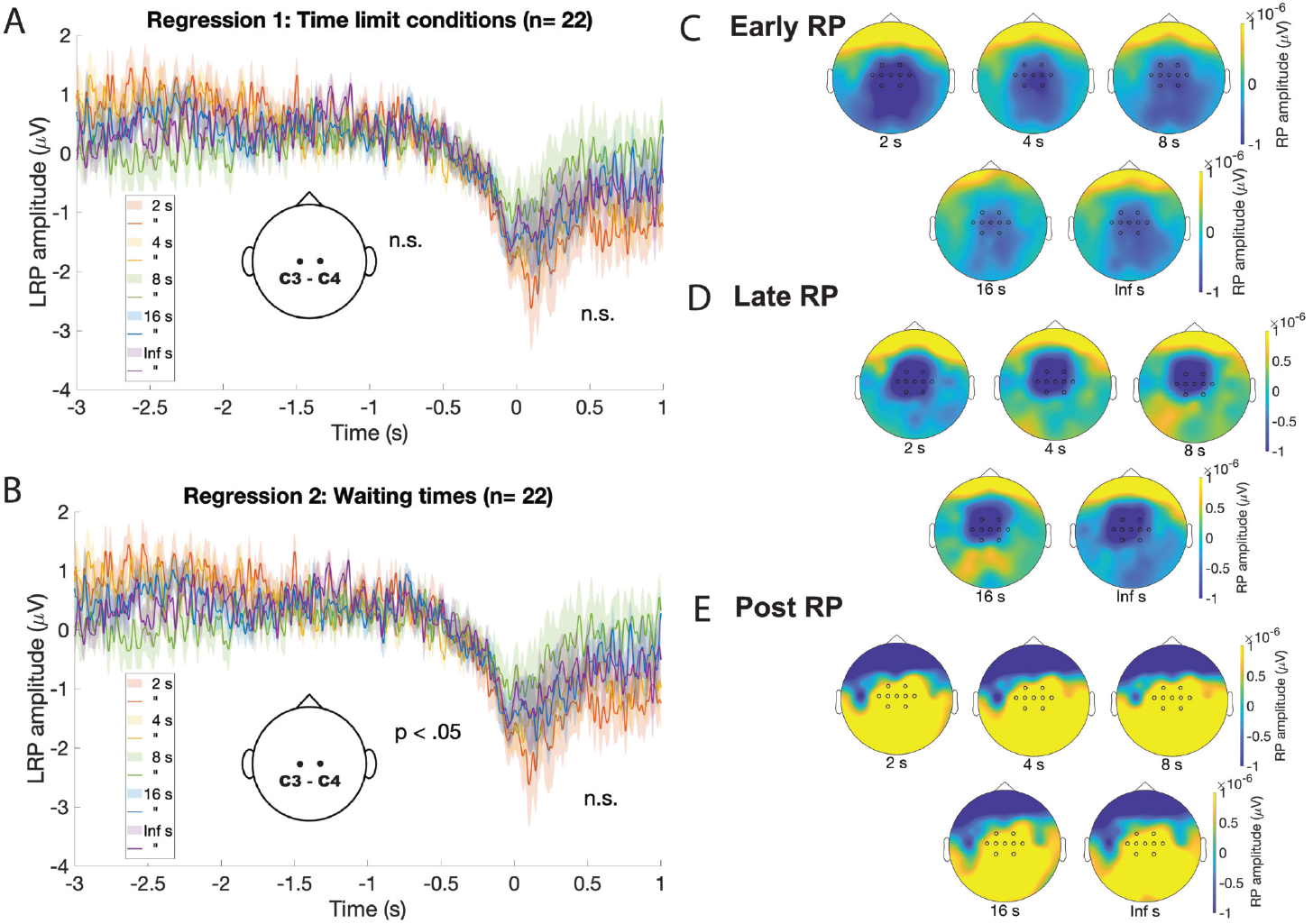
**(A, B)** Grand-averaged LRP amplitude SEM across participants for the conditions 2*s* (red line and shade), 4*s* (yellow line and shade), 8*s* (green line and shade), 16*s* (blue line and shade), *Inf* (violet line and shade). Data are time-locked to time 0 s (finger lift) and result from the grand-average of the subtraction between the EEG signal from electrodes C3 and C4 (LRP= C3 - C4). There were no significant time points (n.s., *p > .*05) resulting from the cluster permutation tests. In **(A)** cluster-based permutation testing was used to test whether the regressions of single-trial LRP amplitudes against the time limit conditions per each trial (predictors in regression 1) was significant. In **(B)** cluster-based permutation testing was used to test whether the regressions of single-trial LRP amplitudes against the single-trial WT responses (predictors in regression 2) was significant. **(C, D, E)** Topography of the grand-averaged RP amplitudes grouped by condition for the latency corresponding to the different phases of the RP, according to the literature: the early RP or pre-commitment phase [−2 s to −0.2 s] **(C)**; the late RP or post-commitment phase [−0.2 s to 0 s] **(D)**, the re-afferent or movement evoked potential [0 s to 1 s] **(E)**. The time limit condition is indicated below each subplot. Note the complete absence of modulation of mean RP amplitude across conditions in the late RP phase as compared to the early RP phase where a clear modulation of mean RP amplitude across conditions is visible.

We sorted the electrophysiological data into two distinct groups, Negative-RPs and Positive-RPs, for group-level statistical analysis and compared it with the Timers and Non-timers subgroups of participants. Given the fact that we had already inspected the data before choosing the final statistical analysis, in order to control the false alarm rate (FA) and avoid biases, we decided to run Fieldtrip’s cluster-based permutation test [42] without pre-defining sensors or time windows of interest. According to the predictions (see Fig. 2) we expected the variables to be positively correlated: an increase in the time limit duration would correspond to an increase in the RP amplitude. However, the data suggested that the variables were instead negatively correlated: an increase in the time limit duration would correspond to a decrease in the RP amplitude, therefore a negative correlation. Since the RP is negative-going by definition, a positive regression between the time limit and the RP magnitude of the signal would mean a more ‘negative’ RP (as it gets farther away from 0) and therefore the appropriate test would be a negative cluster test. A negative regression between the time limit and the RP magnitude of the signal would mean a more ‘positive’ RP (as it gets farther away from 0) and therefore the appropriate test would be a positive cluster test.

### Increasing Time Pressure Induces a Corresponding Decrease in Early RP Amplitudes

For the first regression, testing across all participants (n= 22) for an effect in the early RP latency (−2 s −0.2 s), the cluster-based permutation test revealed a significant positive cluster effect (*p* = 0.0400, cluster based permutation test, two-sided, corrected), but not a significant negative one (*p* = 0.3786, cluster based permutation test, two-sided, corrected), see 4a. Since we had an a priori hypothesis that RP activity should be negative, we performed a two-tailed test on the subgroup of Negative-RPs. Testing across the participants with a clear RP profile or Negative RPs (n= 12) for an effect in the early RP latency (−2 sec −0.2 s), the cluster-based permutation test revealed a significant positive cluster effect (*p* = 0.0300, cluster-based permutation test, two-sided, corrected), but not a significant negative one (*p* = 0.4585, cluster based permutation test, two-sided, corrected), see Table S4. Given the effect found overall across all participants, we further tested across the participants without an identifiable RP profile or Positive-RPs (n= 10) for an effect in the early RP latency (−2 s −0.2 s), the cluster-based permutation test did not find any significant positive cluster (*p* = 0.1229, cluster-based permutation test, two-sided, corrected) or negative cluster (*p* = 0.4835, cluster-based permutation test, two-sided, corrected), see Table S4. Given the relationship between the electrophysiological and the behavioral subgroups, we further performed the cluster-based permutation on the subgroups of Timers and Non-timers. Surprisingly, testing across all the Timers (n= 15) for an effect in the early RP latency (−2 s −0.2 s), the cluster-based permutation test did not find any significant positive cluster (*p* = 0.2148, cluster-based permutation test, two-sided, corrected) or negative cluster (*p* = 0.6224, cluster-based permutation test, two-sided, corrected). Finally, testing across all the Non-timers (n= 7) for an effect in the early RP latency (−2 s −0.2 s), the cluster-based permutation test revealed a significant positive cluster (*p* = 9.9900*e* − 04, cluster-based permutation test, two-sided, corrected) but no a negative one (*p* = 0.1938 cluster-based permutation test, two-sided, corrected).

### Increasing Waiting Times Relates to a Corresponding Decrease in Early RP Amplitudes

For the second regression, testing across all participants (n= 22) for an effect in the early RP latency (−2 s −0.2 s), the cluster-based permutation test revealed a significant positive cluster effect (*p* = 0.0270, cluster based permutation test, two-sided, corrected), but not a significant negative one (*p* = 0.4995, cluster based permutation test, two-sided, corrected), see 4a. Since we had an a priori hypothesis that RP activity should be negative, we performed a two-tailed test on the subgroup of Negative-RPs. Testing across the participants with a clear RP profile (n= 12) for an effect in the early RP latency (−2 sec −0.2 s), the cluster-based permutation test revealed a significant positive cluster effect (*p* = 0.0060, cluster-based permutation test, two-sided, corrected), but not a significant negative one (*p* = 0.2268, cluster based permutation test, two-sided, corrected), see Table S4. Given the effect found overall across all participants, we further tested across the participants without an identifiable RP profile (n= 10) for an effect in the early RP latency (−2 s −0.2 s), the cluster-based permutation test did not find any significant positive cluster (*p* = 0.0549, cluster-based permutation test, two-sided, corrected) or negative cluster (*p* = 0.3576, cluster-based permutation test, two-sided, corrected), see Table S4. Given the relationship between the electrophysiological and the behavioral subgroups, we further performed the cluster-based permutation on the subgroups of Timers and Non-timers. Surprisingly, testing across all the Timers (n= 15) for an effect in the early RP latency (−2 s −0.2 s), the cluster-based permutation test revealed a significant positive cluster (*p* = 0.0030, cluster-based permutation test, two-sided, corrected) but did not find any negative cluster (*p* = 0.3477, cluster-based permutation test, two-sided, corrected). Finally, testing across all the Non-timers (n= 7) for an effect in the early RP latency (−2 s −0.2 s), the cluster-based permutation test revealed a significant positive cluster (*p* = 0.07899, cluster-based permutation test, two-sided, corrected) but no a negative one (*p* = 0.5554 cluster-based permutation test, two-sided, corrected).

### Manipulating the Temporal Pressure Does not Affect Late RP or Lateralized Readiness Potential Amplitudes

Our a-priori hypothesis (see Introduction) concerned the early RP latency (pre-commitment phase), so the window of interest for performing the statistical tests was restricted to −2 s −0.2 s. As a control we tested also for the following temporal window: −0.2 s to 1 s, roughly corresponding to the latency of the late RP starting with the point of no-return (post-commitment phase) [46], the motor evoked potential latency or MEP (which covers the last 50 ms before movement onset) and the post-movement positive complex or re-afferent potential [34]. By regressing the RP amplitudes at each time point against the time limit conditions (first regression) testing for an effect after the point of no-return in all the participants (n= 22), the cluster-based permutation test did not find any significant positive cluster (*p* = 0.3237, cluster based permutation test, two-sided, corrected), or negative cluster (*p* = 0.1668, cluster based permutation test, two-sided, corrected), see 4a. When testing for an effect in the same latency in the Positive-RPs participants (n= 10), the cluster-based permutation test did not find any significant positive cluster (*p* = 0.3027, cluster based permutation test, two-sided, corrected), however it did find a significant negative cluster (*p* = 0.0340, cluster based permutation test, two-sided, corrected), see Table S4. When testing for an effect after the point of no-return in the Timers participants (n= 15), the cluster-based permutation test did not find any significant positive cluster (*p* = 0.5874, cluster based permutation test, two-sided, corrected) or significant negative cluster (*p* = 0.5355, cluster based permutation test, two-sided, corrected), see Table S4. When we regressed the RP amplitudes at each time point against the behavioral waiting time responses for the same window of interest (−0.2 s to 1 s) in all the participants (n= 22), the cluster-based permutation test did not find any significant positive cluster (*p* = 0.2128, cluster based permutation test, two-sided, corrected), or negative cluster (*p* = 0.4216, cluster based permutation test, two-sided, corrected), see 4b. When testing for an effect in the Negative-RPs participants (n= 12), the cluster-based permutation test did not find any significant positive cluster (*p* = 0.3017, cluster based permutation test, two-sided, corrected), or negative cluster (*p* = 0.8322, cluster based permutation test, two-sided, corrected), see Table S4. When testing for an effect in the Positive-RPs participants (n= 10), the cluster-based permutation test did not find any significant positive cluster (*p* = 0.4835, cluster based permutation test, two-sided, corrected) nor a significant negative cluster (*p* = 0.2507, cluster based permutation test, two-sided, corrected). When testing for an effect in the Timers participants (n= 15), the cluster-based permutation test did not find any significant positive cluster (*p* = 0.6513, cluster based permutation test, two-sided, corrected) or significant negative cluster (*p* = 0.8342, cluster based permutation test, two-sided, corrected), see Table S4. When testing for an effect in the Non-timers participants (n= 7), the cluster-based permutation test did not find any significant positive cluster (*p* = 0.1049, cluster based permutation test, two-sided, corrected) but it did find a significant negative cluster (*p* = 9.9900*e* − 04, cluster based permutation test, two-sided, corrected), see Table S4.

Finally, an important control was the presence of a Lateralized Readiness Potential (LRP), see 5, which confirmed the lateralization of the response preparation, expected since in our instruction to the participants we had specified the response side (right hand for the finger lift). To compute the LRP we subtracted the mean amplitude of sensors C3 and C4 (see Methods). We tested for parametric effects in the LRP through regression of the LRP amplitudes at each time point against either the time limit conditions or the behavioral waiting time responses and as a control we used the same latencies and subgroups of subjects as for the RP. In the first regression we found no significant positive or negative clusters (*p > .*05, cluster-based permutation test, two-side, corrected) in the latency −2 s −0.2 s for neither all subjects together (n= 22), see 5a, Negative RPs (n= 12), Positive RPs (n= 10), Timers (n= 15), NonTimers (n= 7), see Table S4. In the latency −0.2 s to 1 s we found no significant positive or negative clusters (*p > .*05, cluster-based permutation test, two-side, corrected) for neither all subjects together (n= 22), see 5b, Positive RPs (n= 10), Timers (n= 15), NonTimers (n= 7). However, we found a significant positive cluster when testing Negative RPs (n= 12) subjects.

Taken together, these results show how a systematic effect of parametrically varying time pressures is selective for the early RP latency and motor preparation phase, and does not concern motor components involved in the movement execution phase.

## Discussion

So far, most studies in the domain of self-initiated movements have focused on the dichotomy between stimulus-triggered and internally-triggered movements [47] without exploring the spectrum in between. Those studies have shown functional and anatomical dissociation between the neural signals preceding self-generated and externally-generated movements both in terms of the signal average and its variability before movement onset [2, 48]. According to a theoretical prediction of the leaky stochastic accumulator model there is not a dichotomy but a continuum from ‘exogenous’ to ‘endogenous’ that can potentially be described in terms of a strong or weak imperative (or drift term). However, it has remained so far experimentally untested how neural activity dynamically evolves along those extremes. Moreover, the current versions of the model (the output and input hypotheses, see [17, 26]) have only tested the possibility of a fixed imperative, for which variability in behavioural responses (giving rise to fast or slow waiting times) depend solely on noise. We wondered if by parametrically manipulating the strength of the imperative to move, operationalised as exogeneous triggers about when to move, we would affect the RP profile and consequently the WT data. In order to do that, we provided participants with a clock displaying parametrically varying levels of time pressures within which they had to make a finger movement: this allowed us to get a closer look at the evolving dynamic of the RP morphology.

At the behavioural level we found that the external imposition of time pressure correlated with participants’ waiting times (meaning that most of the participants waited longer if there was more time allotted). At the electrophysiological level the time pressure-imperative to move anti-correlated with the time-locked RP signal amplitude (the EEG signal was ‘stronger’ the shorter the limits of time to perform a movement). However, this effect was specific to the early phase of the RP up to about −200 ms before movement onset: neither the very late portion of the RP or the post-motor positivity (PMP) exhibited a change in amplitude corresponding to the experimental manipulation. To further corroborate this finding, as a control, we also tested for the presence of a significant modulation induced by the task in the amplitude of the lateralized readiness potential (LRP) [40, 49] without finding any.

Traditionally, the RP is divided into two physiologically and functionally distinct components: the early RP, which originates bilaterally in the pre-supplementary motor area (pre-SMA), and is known to be influenced by cognitive factors such as “level of intention, preparatory state and movement selection”; the late RP, whose generator source is lateralized in the contralateral primary motor cortex (M1) and lateral pre-motor cortex, and is supposedly affected by motor related features such as precision, discreteness and complexity of the movement [34]. This distinction is consistent with that between central processes responsible for programming the movement and peripheral processes implicated in the initiation of the movements, as dual-process motor theories posit [50]. Schurger et al.’s model [17] interprets, instead, the motor preparation phase as the input evidence preceding the decision to move now, a neural commitment represented by the threshold crossing moment (at around 150 ms), and, consequently, the motor execution phase as the output decision variable following this threshold crossing event. Therefore, the pre-commitment phase or early RP is dominated by stochastic fluctuations, while the post-commitment phase or late RP corresponds to a lateralization of the readiness potential in an effector-dependent manner [51] and an increase in cortical excitability [52]. This theoretical framework is compatible with two recent experimental findings. Concerning the post-commitment phase, Schultze-Kraft et al. found a point of non return (at around −200 ms) [46] after which a motor signal progressing to the peripheral muscles in a ballistic way [53] cannot be cancelled any longer once the brain enters the motor execution stage. Concerning the pre-commitment phase, Khalighinejad et al. [48] found that the across-trial decrease in variability of the stochastic fluctuations before self-initiated movements is stronger in fronto-midline electrodes (like Cz) which are associated with a general cognitive preparation process independent of the hand that will execute the movement.

Our results converge with these studies in showing at around −200ms this transition from a regime governed by a neural decision process, affected by external triggers or endogenous activity, to one where the decision unfolds into an actual, ballistic movement. It has to be noticed that in [17] the authors arbitrarily choose to fit the stochastic decision model output to the RP signal up to −150 ms, based on previous literature. However, the present study brings the first experimental evidence validating the model’s assumptions regarding the early and late RP.

So far we interpreted the data in terms of a manipulation of the imperative to move, however another interpretation is possible: in the very first RP study [14, 15] the readiness potential is shown to increase “with intentional engagement” and be reduced “by mental indifference of the subject”. A follow up study by McAdam and Seale (1969) [54] confirmed an enhancement in the RP amplitude due to increasing levels of motivation. Freude and Ullsperger (1988) [55] showed that the RP is significantly higher in amplitude when participants had to solve tasks involving mental load under higher time pressure, even with motor activity kept constant. If we consider the fact that in our experiment participants are trained to make a finger movement specifically within the allotted window of time and are negatively reinforced for being too early or too late on the response, we cannot rule out the possibility that the observed effect on the RP is instead a consequence of the task demands. Indeed, it might be that for shorter time limits, where the pressure to make a response is higher, participants feel more ‘engaged’ in their movement, while in the longer time limits, where they are free to wait as much as they want, participants feel more ‘relaxed’. Along the same line that links the RP to non motoric mechanisms, Pornpattananangkul and Nusslock (2015) [56] adapted a reward time-estimation task in which participants were instructed to make a button-press 3.5 s after the onset of a cue and were rewarded for accurate estimation. They showed that the RP amplitude significantly increased for the trials belonging to the reward condition. Alexander et al. (2016) [57] claimed that RPs can occur in absence of movements as their amplitude is modulated by decision and anticipation. More recently, Wen et al. (2018) [58] argued that the awareness of action-effect contingency can have an internally rewarding effect for the action itself and therefore increase the RP amplitude.

It could be that the current design is manipulating something other than the imperative to move, such as the above-mentioned cognitive processes or temporal expectation. For example someone could argue that some components of the task resembled the contingent negative variation (CNV) paradigm [59]: the visual cue might resemble the warning stimulus and the time limit target bar the imperative stimulus. Therefore the cortical negativity that we captured through epoching and time-locking could be just a portion of a CNV curve. However, in classic CNV tasks the expectation is generated by the association between a fixed warning and an imperative cue. In our case, to prevent subjects from building up a prediction about when to move we included random times both at the starting of the moving dot and after the time-limit was passed (the inter-trial interval).

In previous studies [17, 26] the stochastic decision model model in the form of a fixed boundary model could provide a satisfactory fitting for both behavioral and electrophysiological data in a self-initiated movement task. However, in the present study model simulations have not managed to reproduce completely the qualitative pattern in the results leaving open the possibility that the current state of the SDM cannot account for the time limit variations paradigm. In perceptual decision-making tasks involving a response deadline participants achieve optimality in performance by collapsing the decision threshold such that the decision is reached on time with less evidence accumulated. Besides the deadline duration, another factor modulating the speed-accuracy trade off is the ‘endogenous timing uncertainty’ [60]: the shorter the deadline duration or the more uncertain the deadline, the earlier the threshold should collapse to allow for a timely response. In our time limit task, the uncertainty about when to move is maximal in the *Inf s* condition, where cues for the duration of the trial are absent, and minimal in the time limit conditions, where a small visual target signaled the deadline for the response. Even if the majority of participants declared in the questionnaire to not have adopted a strategy, the tendency was to not wait longer than a quarter of the time allotted. In conditions of equal endogenous uncertainty for performing the response, the differences among the waiting times might have reflected a time-dependent threshold decay, for which a collapsing boundary model provides a better description. Indeed, participants on average waited less for the *Inf s* condition (longest deadline but least certain) rather than for the *16 s* the condition (long deadline but more certain). A previous study (Gluth et al. 2013) [61] successfully modeled the influence of evidence accumulation and elapsed time on the RP with a time-variant sequential sampling model (SSM) that assumes threshold collapse. The need for such models has been deemed controversial for being redundant with the standard diffusion model and not providing an improvement for tasks with human data [62, 63]. However, a recent line of research has proposed a shift from conventionally behavior-based drift-diffusion models (DDM) with one source of build-up (evidence accumulated) to neurally informed (NI) decision models with multiple build-up (evidence accumulated and dynamic, evidence-independent urgency signal) and time delay components [18, 64, 65]. In particular, Kelly et al. (2020) have experimentally shown a better fit for neurophysiological data through the NI model as compared to the classic DDM. Moreover, in convergence with our empirical results and relative to a speed pressure paradigm, the authors have found an enhancement of the drift rate under short deadlines (‘deadline regime’) in contrast to long deadlines (‘easy regime’), whereas the DDM fit lead to opposite conclusions analogous to the qualitative predictions of the SDM which was also constructed upon behavioural data. Further work should try to address these issues by proposing a more complex version of the model that fully characterizes the dynamic aspects of the task (such as different levels of time-dependent bounds, urgency signal) by leveraging neural data to quantitatively constrain the model parameters (see also [66]).

An established fact in the readiness potential literature is the fact that the signal precedes only self-initiated movements and not stimulus-triggered ones. This makes sense from the perspective of the classical view which entails a preparatory function for the RP shape. However, according to the stochastic model, the RP is originated by averaging ongoing fluctuations time-locked to movement that actually occurred.

Therefore, the RP is not ‘exclusive’ to internally initiated movements but can be originated also in externally triggered cases. Indeed, readiness potentials could be recovered in the average of reaction time trials in a task in which movement that was not intended to be performed was induced by unpredictable, compulsive auditory cues (the Libetus interruptus task, [17]). Here, we could replicate this finding by obtaining RPs also when the movement timing was not completely freely decided because of the time constraint.

A pillar regarding the nature of the RP is its slowly-increasing negative voltage [23, 34, 67, 68]. However, one of the findings of our study was that not all participants showed the same polarity in the RP, with 45% of them displaying a positive shift and some presenting no RP at all. This is not a novel account though it has received minor attention in the RP literature: Freude and Ullsperger (1989) [69] had already found in single-trial analysis that 44% of the RPs preceding self-paced voluntary movements present slow positive potentials shifts; VaezMousavi and Barry (1993) [70] replicated the same results in another experiment and showed that the proportion of positive RPs increased by 10% in conditions of mental load, suggesting a relationship between the effects of spontaneous brain activity on improved cognitive performance and the negative RP polarity. More recently, another group (Jo et al., 2013) [71] reported that the apparent negative slope in the average RP results as the sum of an unequal ratio of negative (67.16%) and positive (32.84%) potential shifts and proposed that the ongoing negative deflections in the slow cortical-potentials (SCPs) facilitate movement initiation by increasing its likelihood (see also the slow cortical potential or SCP sampling hypothesis [72]). In our study, however, we report the presence of positive shifts not on single-trial level but on averaged trials within and across-subjects. Therefore, contrary to previous records for which positive RPs are ‘cancelled out’ in a weighted average with more frequent negative RPs, we show for the first time RP grand-averages with an overall positive dominant polarity, indicating a bigger proportion of positive single-trial epochs in comparison to the negative ones. Surprisingly, including subjects with a prevalent positive RP profile in the statistical analysis did not make the results less significant. This result is unexpected and suggests that the effect due to the parametric variation of the imperative to move persists regardless of the polarity of the RP. It is worth noting that the stochastic decision model predictions are agnostic regarding the sign of the RP voltage, though this is usually assumed to be negative.

Another unprecedented finding related to the RP polarity is the striking overlap between the subjects grouped according to their RP polarity (‘positive RPs’ or ‘negative RPs’) and the same subjects grouped according to their individual variability in their waiting time responses across different conditions (‘timers’ or ‘non-timers’). If the RP negativity reflects increased cortical excitability correlated with motor performance, as suggested by the SCP theory [73], we could speculate that this is the reason why the participants whose behavioural responses followed task demands coincide with those with the predominant negative shifts. Further analyses will be needed to test for this hypothesis.

Overall, these findings about how the shape of the RP varies, across experimental conditions (the imperatives to move), how its polarity changes, across and even within subject, and how the RP is generated suggest that the RP is not a highly stereotypical signal as commonly believed [46].

## Conclusion

In conclusion, we developed a novel paradigm that extends the traditional Libet task to a continuum spanning from temporal pressure to temporal freedom which corresponded to more stimuli-driven decisions to move and internally generated decisions to move. We showed that the parametric variation of temporal pressures affects movement related neural activity. We found that increasing the time windows allotted for making a self-initiated movement was associated with a decrease in the readiness potential’s amplitude corresponding to its early components. We also showed that the manipulation did not affect either the late component of the RP (after the so called ‘point of no-return’) or the LRP. Finally, we suggest a possible revision of the stochastic decision model with the inclusion of collapsing bounds with different rates of collapse or dynamic urgency components that could provide a more neurally plausible account for time-limit or deadline type of tasks.

## Supporting information

**File S1. Questionnaire in French.** We report the original anonymous questionnaire sheet delivered to the participants at the end of the experiment. Individual responses are available upon request.

**Fig. S1. Behavioural results. (A, B)** Distribution of subjects waiting times sorted by time-limit condition 2*s* (red), 4*s* (yellow), 8*s* (green), 16*s* (blue), *Infs* (violet). Note that the shape of the distribution of the WTs is very similar across conditions no matter the time-limit duration (time-scale invariance). **(C)** Subjects standard deviations of the WTs are sorted by condition. Subjects whose WTs standard deviations do not scale following the order of the time-limit conditions are framed. **(D)** Linear relationship between the mean and the standard deviation of WTs, mechanism typical from perceptual-decision making task.

**Fig. S2. Individual RPs profiles. (A)** Averaged RP amplitude within participants (n= 22) ± SD for the condition *Infs* (violet line and shade) over 4 s (epoch: −3 s to 1 s). Data are time-locked to time 0 s (finger lift). For representative purposes, we display EEG activity recorded from Cz electrode which did not differ from the ROI cluster of channels. Red lines indicate slopes computed over the last 1 s before movement onset. **(B)** Topography of the grand-averaged RP amplitudes within participants (n= 22) for the condition *Infs* and the latency corresponding to the last 1 s before movement onset. All topographic plots have been normalized to the same ERP amplitude scale. ROI channels are highlighted.

**Fig. S3. Negative-RPs, Positive-RPs, Timers, Non-timers. (A)** Averaged RP amplitude ± SEM across participants displaying a canonical, negative RP (n= 12) and bar plots representing averaged RP amplitudes across conditions from −2 s to 0 s for the same subgroup of subjects. **(B)** Grand-averaged RP amplitude ± SEM across participants displaying a positive or flat RP (n= 10) and bar plots representing averaged RP amplitudes across conditions from −2 s to 0 s for the same subgroup of subjects. **(C)** Grand-averaged RP amplitude ± SEM across participants whose std WTs did not scale with the time-limit duration (n= 15) and bar plots representing averaged RP amplitudes across conditions from −2 s to 0 s for the same subgroup of subjects. **(D)** Grand-averaged RP amplitude ± SEM across participants whose std WTs did not scale with the time-limit duration (n= 15) and bar plots representing averaged RP amplitudes across conditions from −2 s to 0 s for the same subgroup of subjects. As in Fig. 3 the conditions and color codes are: 2*s* (red line and shade), 4*s* (yellow line and shade), 8*s* (green line and shade), 16*s* (blue line and shade), *Inf* (violet line and shade). Data are time-locked to time 0 s (finger lift). For representative purposes, we display EEG activity recorded from Cz electrode which did not differ from the ROI cluster of channels.

**Table S1. Results from one-sample Kolgoromov-Smirnov test.** We run post-hoc pair-wise comparisons comparing the behavioural responses in each condition to all the others conditions in order to reveal where the effect of the time limit conditions were.

**Table S2. Results from the post-hoc multiple comparison test.** We computed one-directional Wilcoxon paired tests between pairs of conditions at *alpha* = 0.05: mean RP amplitudes and slopes within each condition for channel FC1, C3, Cz separately and ROI (average across channels) and mean RP amplitudes obtained through the EMSF technique. The uncorrected p-values were all adjusted with the Benjamini and Hochberg (1995) procedure for controlling the false discovery rate (FDR) for multiple comparison correction. (*) indicates p-values *<*0.05.

**Table S3. Cluster-based permutation tests results.** We report the results of the nonparametric statistical tests performed with the method Monte-Carlo on the beta coefficients of the regressions. *cfg.clusteralpha*, *cfg.correcttail*, *cfg.neighbour* correspond to Fieldtrip parameters. *stat.posclusters.prob* is the output of the permutation test for positive clusters. (*) indicates p-values <0.05. (**) indicates p-values <0.005.

## Author Contributions

<u>Bianca Trovò</u>: Conceptualization, Methodology, Software, Validation, Formal analysis, Investigation, Resources, Data Curation, Writing - Original Draft, Writing - Review & Editing, Visualization, Project administration; <u>Yvonne Visser</u>: Investigation, Project Administration, Validation, Writing - Review & Editing; <u>Zafer Isčan</u>: Software, Writing - Review & Editing; <u>Aaron Schurger</u>: Conceptualization, Funding acquisition, Methodology, Software, Writing - Review & Editing, Visualization. All authors approved the submitted version.

## Acknowledgments

We thank Virginie van Wassenhove, Sophie Herbst, Florent Meyniel, Maxime Mahieu for scientific advice and discussions on the EEG analysis and statistics; Giulia Gennari for feedback on the figures and comments on the manuscript. We thank the Neurospin nurses (LBIOM) and the UNIACT staff for their help in recruiting participants for the experiment.

## Notes

### Competing Interest Statement

The authors have declared no competing interest.

