## Supplemental File 1 for "Movement-Preceding Neural Activity under Parametrically Varying Levels of Time Pressure"

### Questionnaire

Experience de limit temporel
Janvier 2018

Participant:

**Est-ce que vous pensez que c'était facile de suivre les instructions sur quand faire le mouvement?**

- Oui ☐ Non ☐

Si non, pourquoi? …………………………………………………………………………

……………………………………………………………………………………………

……………………………………………………………………………………………

**Est-ce que votre action était spontanée à chaque essai?**

- Toujours ☐ Plus part du temps ☐ Parfois ☐ Presque jamais ☐ Jamais ☐

Si pas toujours, pourquoi? …..…………………………………………………………

……………………………………………………………………………………………

……………………………………………………………………………………………

**Est-ce que vous avez utilisé une stratégie pour déterminer quand bouger?**

……………………………………………………………………………………………

……………………………………………………………………………………………

……………………………………………………………………………………………

**Est-ce que vous pensez qu'un facteur externe puisse avoir influencé votre décision sur quand faire le mouvement?**

……………………………………………………………………………………………

……………………………………………………………………………………………

……………………………………………………………………………………………

**Est-ce que vous pouvez décrire votre expérience?**

……………………………………………………………………………………………

……………………………………………………………………………………………

……………………………………………………………………………………………

……………………………………………………………………………………………
