## Supplemental Figure 1 for "Movement-Preceding Neural Activity under Parametrically Varying Levels of Time Pressure"

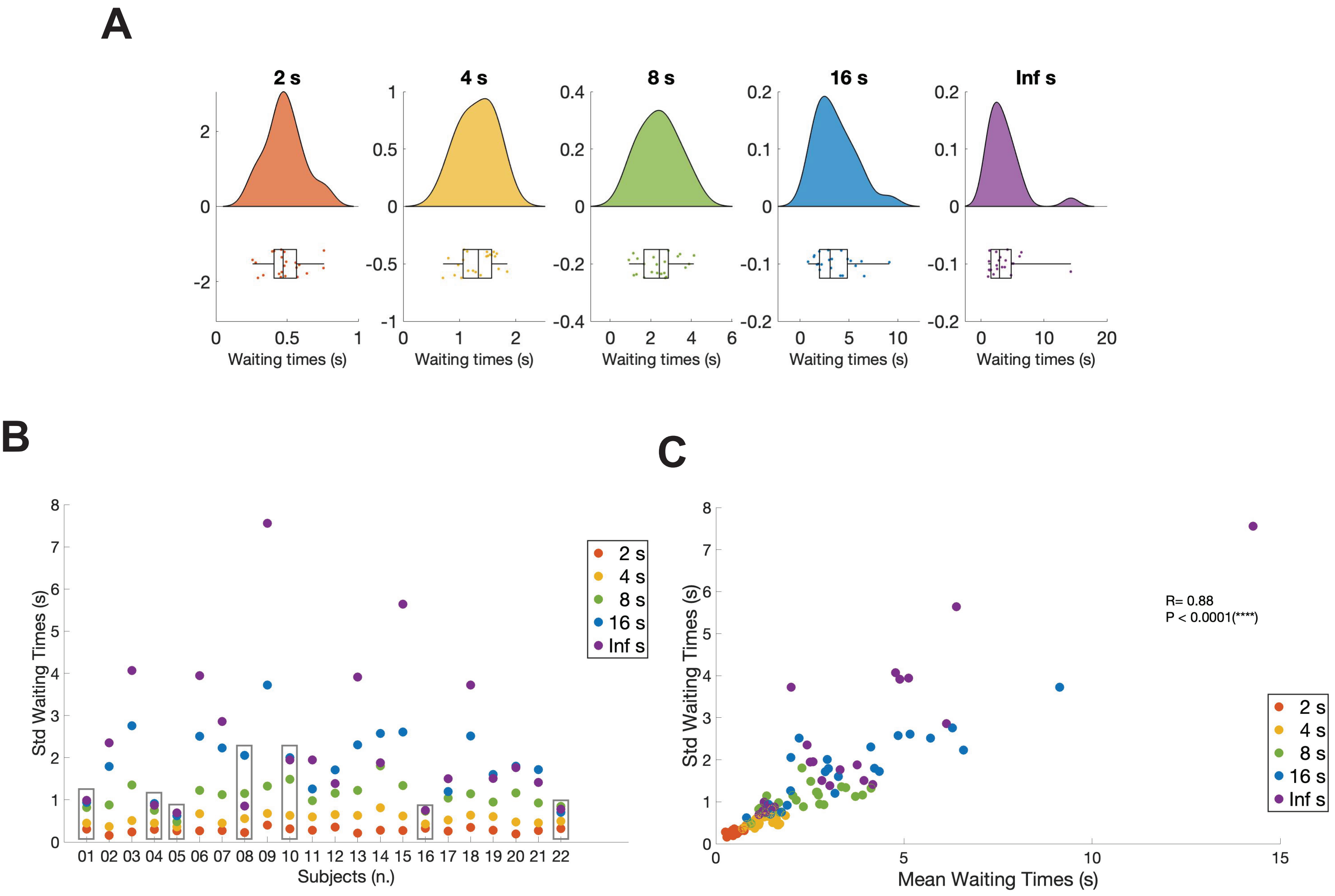


**Fig. S1. Behavioural results. (A)** Distribution of subjects waiting times-sorted by time-limit condition 2 s (red), 4 s (yellow), 8 s (green), 16 s (blue),Inf s (violet). Note that the shape of the distribution of the WTs is very similar across conditions no matter the time-limit duration (time-scale invariance). **(B)** Subjects standard deviations of the WTs are sorted by condition. Subjects whose WTs standard deviations do not scale following the order of the time-limit conditions are framed. **(C)** Linear relationship between the mean and the standard deviation of WTs, mechanism typical from perceptual-decision making task.
