## Supplemental Figure 2 for "Movement-Preceding Neural Activity under Parametrically Varying Levels of Time Pressure"

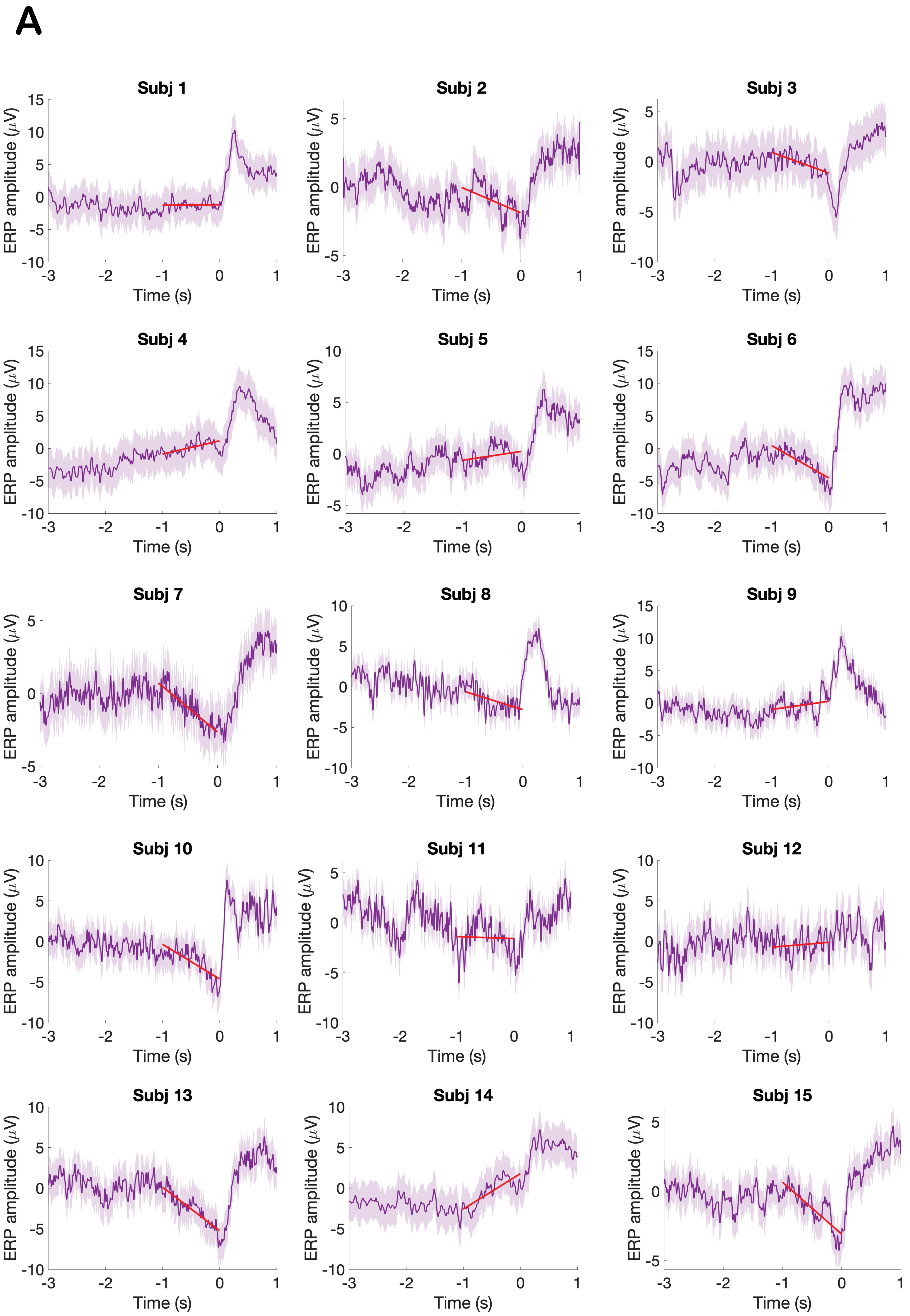
A
Subj 1
Subj 2
Subj 3
Subj 4
Subj 5
Subj 6
Subj 7
Subj 8
Subj 9
Subj 10
Subj 11
Subj 12
Subj 13
Subj 14
Subj 15
ERP amplitude ($\mu$V)
Time (s)

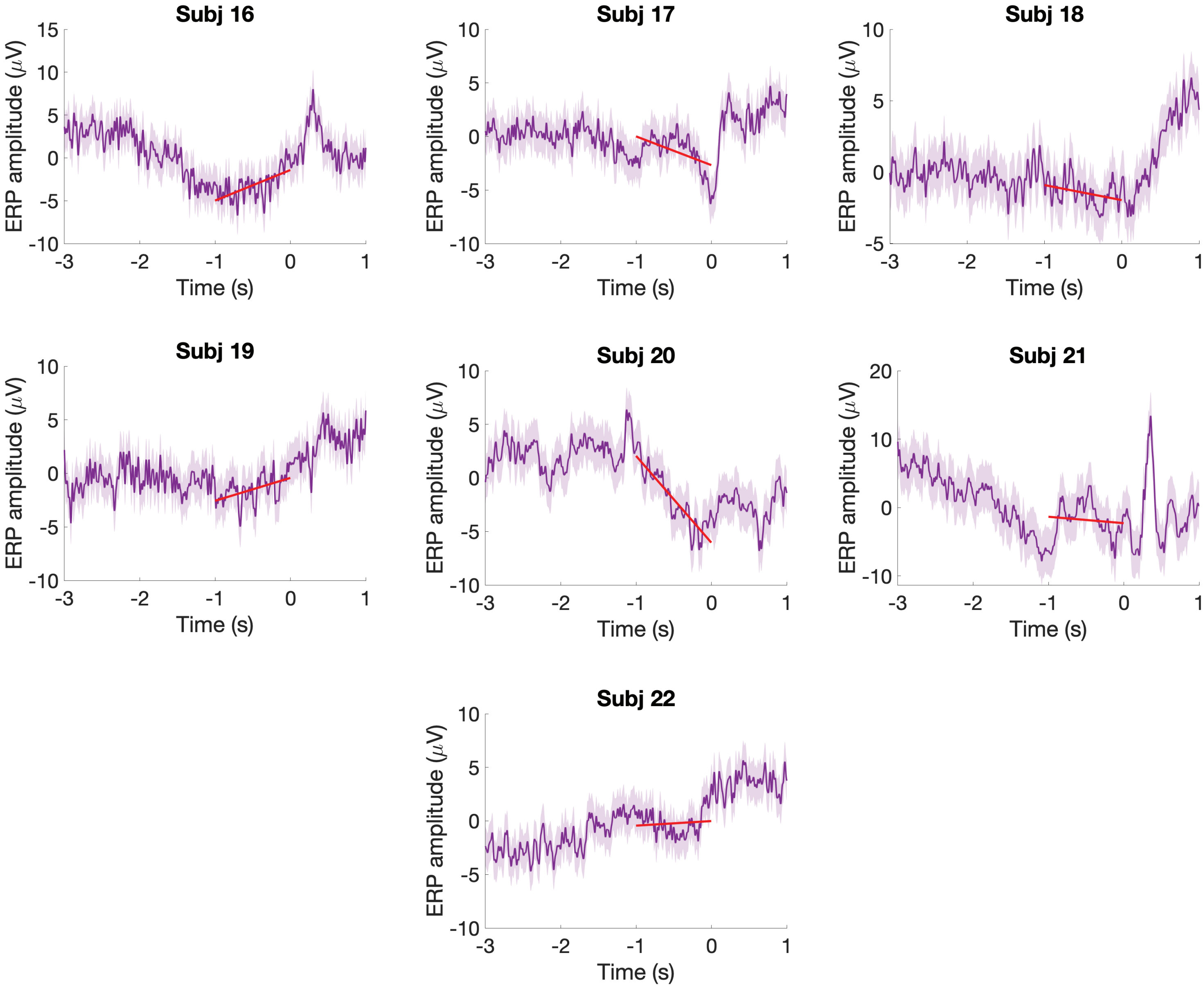

**Fig. S2. Individual RPs. (A)** Averaged RP amplitude within participants (n= 22) ± SD for the condition Inf s (violet line and shade) over 4 s (epoch: -3 s to 1 s). Data are time-locked to time 0 s (finger lift). For representative purposes, we display EEG activity recorded from Cz electrode which did not differ from the ROI cluster of channels. Red lines indicate slopes computed over the last 1 s before movement onset. 12 subjects (3, 6, 7, 8, 10, 13, 15, 17, 18, 19, 20, 21) were classified as Negative-RPs because exhibit a stereotypical RP in the Inf s condition. 10 subjects (1, 2, 4, 5, 9, 11, 12, 14, 16, 22) were classified as Positive-RPs because the RP could not be identified (positive-going or flat RP signal).

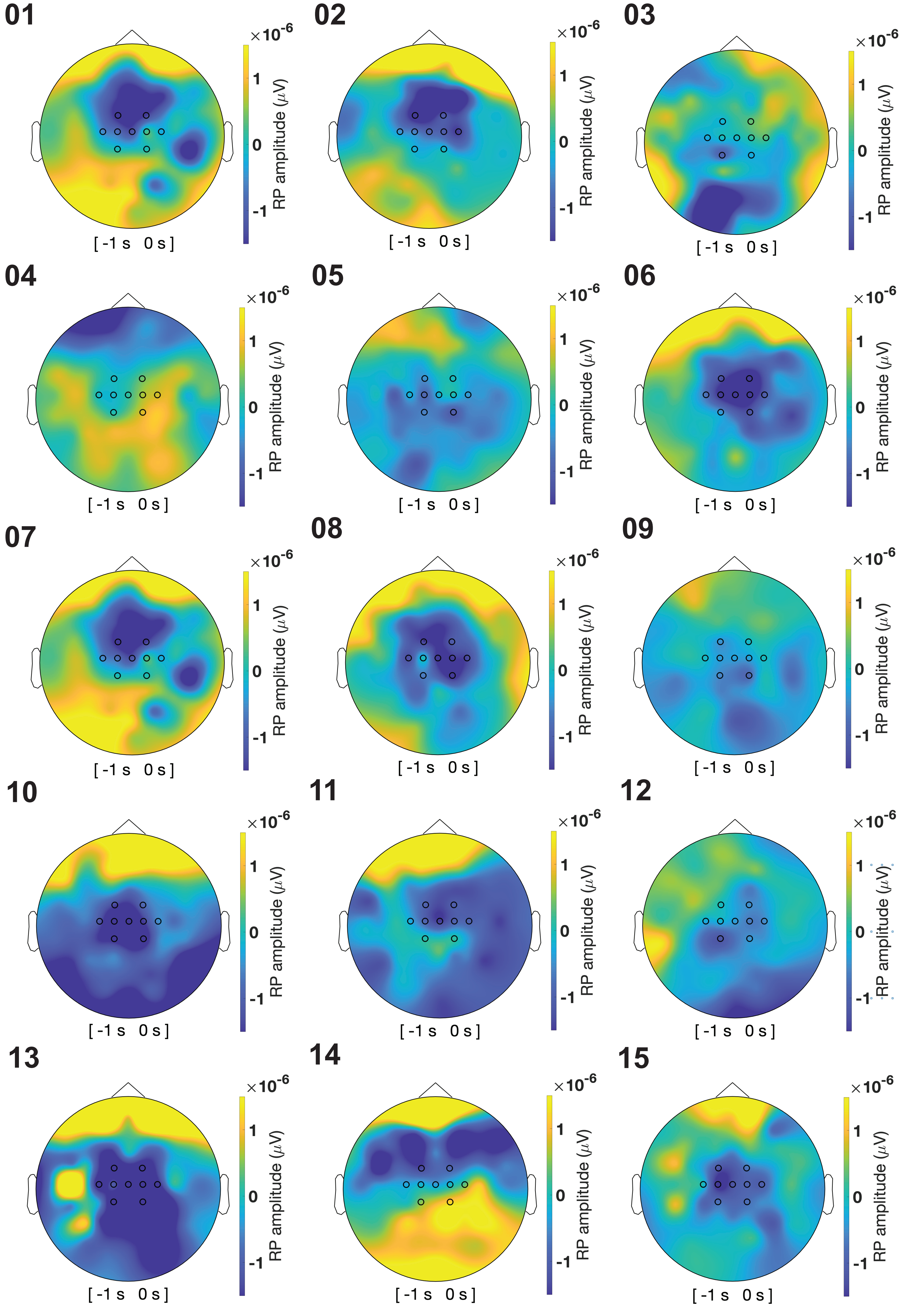

×10^-6
1
0
-1
RP amplitude (μV)
[-1 s 0 s]

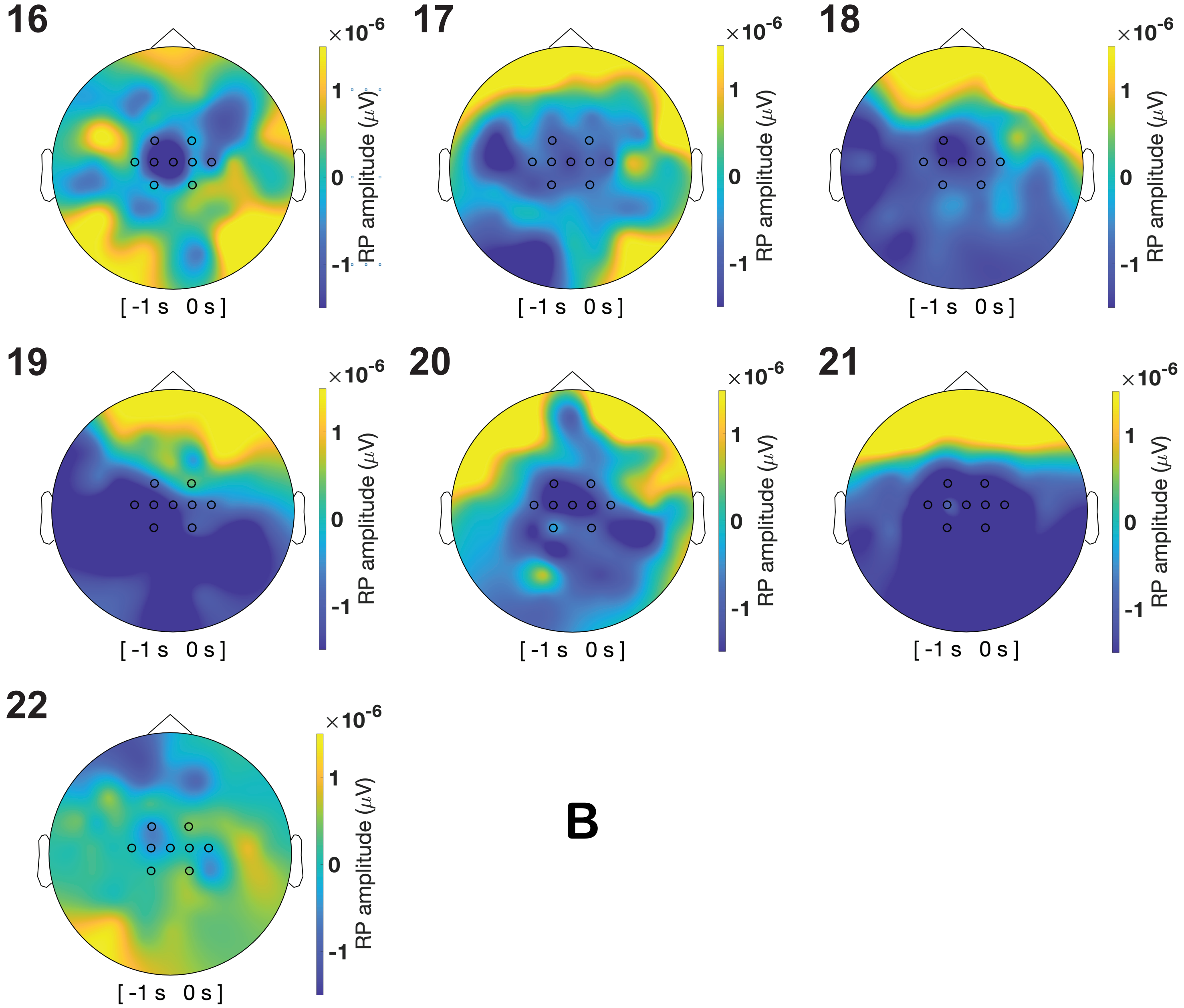

**Fig. S2. Individual RPs. (B)** Topography of the averaged RP amplitudes within participants (n= 22) for the condition Inf s and the latency corresponding to the last 1 s before movement onset. All topographic plots have been normalized to the same RP amplitude scale. ROI channels are highlighted. 12 subjects (3, 6, 7, 8, 10, 13, 15, 17, 18, 19, 20, 21) were classified as Negative-RPs because exhibit a stereotypical RP in the Inf s condition (negativity spreading from the center of the scalp). 10 subjects (1, 2, 4, 5, 9, 11, 12, 14, 16, 22) were classified as Positive-RPs because the RP could not be clearly identified.
