## Supplemental Figure 3 for "Movement-Preceding Neural Activity under Parametrically Varying Levels of Time Pressure"

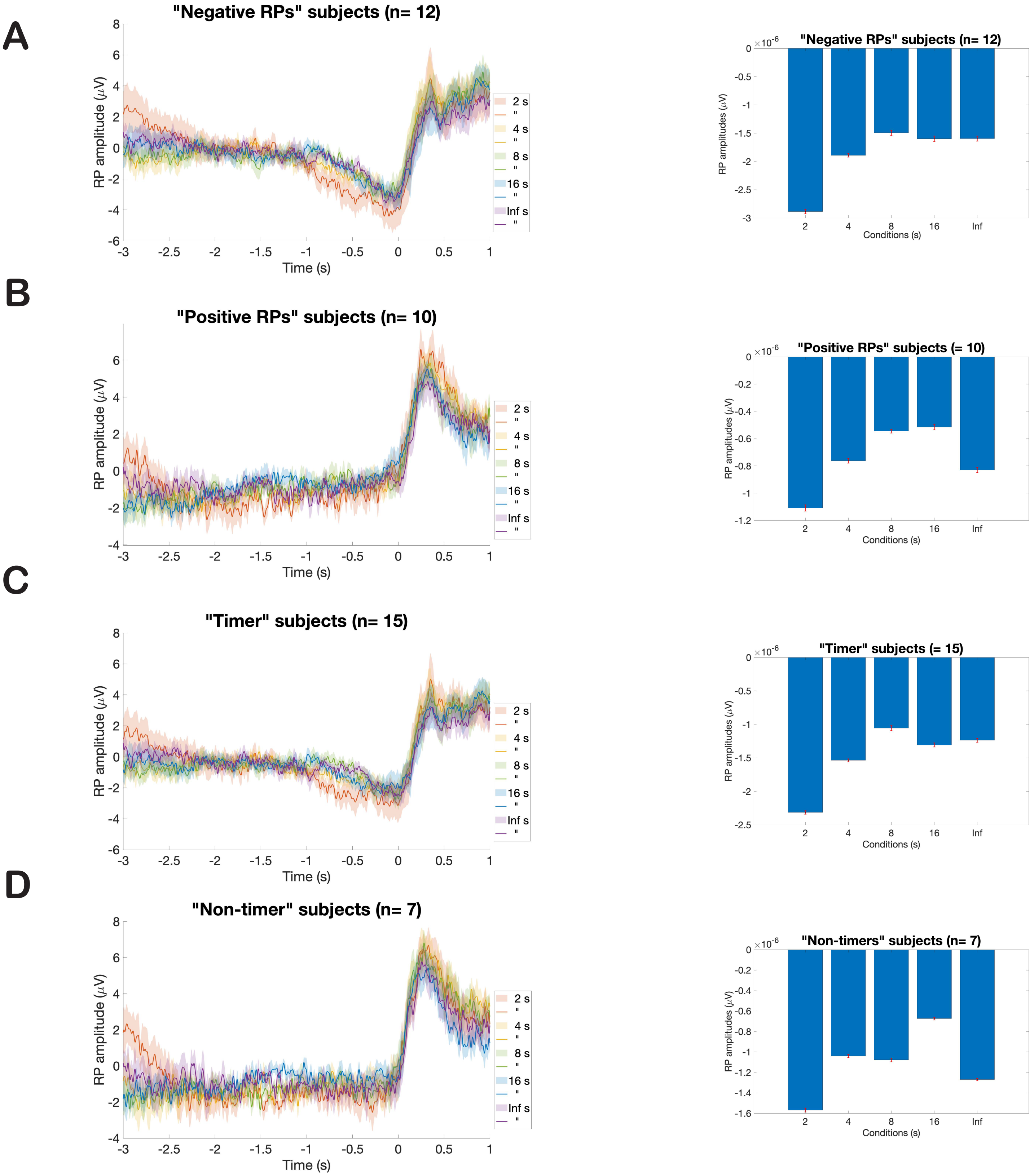


**Fig. S3. Negative-RPs, Positive-RPs, Timers, Non-timers. (A)** Grand-averaged RP amplitude ± SEM across participants displaying a canonical, negative RP (n= 12) and bar plots representing averaged RP amplitudes across conditions from -2 s to 0 s for the same subgroup of subjects. **(B)** Grand-averaged RP amplitude ± SEM across participants displaying a positive or flat RP (n= 10) and bar plots representing averaged RP amplitudes across conditions from -2 s to 0 s for the same subgroup of subjects. **(C)** Grand-averaged RP amplitude ± SEM across participants whose std WTs did not scale with the time-limit duration (n= 15) and bar plots representing averaged RP amplitudes across conditions from -2 s to 0 s for the same subgroup of subjects. **(D)** Grand-averaged RP amplitude ± SEM across participants whose std WTs did not scale with the time-limit duration (n= 15) and bar plots representing averaged RP amplitudes across conditions from -2 s to 0 s for the same subgroup of subjects. As in Fig. 3 the conditions and color codes are: 2 s (red line and shade), 4 s (yellow line and shade), 8 s (green line and shade), 16 s (blue line and shade), Inf (violet line and shade). Data are time-locked to time 0 s (finger lift). For representative purposes, we display EEG activity recorded from Cz electrode which did not differ from the ROI cluster of channels.
