## Supplemental Table 1 for "Movement-Preceding Neural Activity under Parametrically Varying Levels of Time Pressure"

**Table S1. Results from one-sample Kolgoromov-Smirnov test.** We tested the normality of the behavioral distribution of the data in each condition. A significant results means that the data is non-normally distributed.

| Condition n° | Statistics | p-values (1.0e-07 *) |
|---|---|---|
| 1 | 0.6014 | 0.0000 |
| 2 | 0.7605 | 0.0000 |
| 3 | 0.8272 | 0.0000 |
| 4 | 0.8744 | 0.0000 |
| 5 | 0.8743 | 0.0000 |

**Table S2. Results from the post-hoc multiple comparison test (Mathworks; Hochberg and Tamhane, 1987).** We run post-hoc pair-wise comparisons comparing the behavioural responses in each condition to all the others conditions in order to reveal where the effect of the time limit conditions were.

| Comparisons | Confidence Interval | | Statistics | p-values |
|---|---|---|---|---|
| 1 vs 2 | − 56.2348 | − 30.0000 | − 3.7652 | 0.0156 |
| 1 vs 3 | − 81.8712 | − 55.6364 | − 29.4015 | 0.0000 |
| 1 vs 4 | − 95.1439 | − 68.9091 | − 42.6743 | 0.0000 |
| 1 vs 5 | − 91.2348 | − 65.0000 | − 38.7652 | 0.0000 |
| 2 vs 3 | − 51.8712 | − 25.6364 | 0.5985 | 0.0592 |
| 2 vs 4 | − 65.1439 | − 38.9091 | − 12.6743 | 0.0005 |
| 2 vs 5 | − 61.2348 | − 35.0000 | − 8.7652 | 0.0025 |
| 3 vs 4 | − 39.5076 | − 13.2727 | 12.9621 | 0.6405 |
| 3 vs 5 | − 35.5985 | − 9.3636 | 16.8712 | 0.8672 |
| 4 vs 5 | − 22.3257 | 3.9091 | 0.1439 | 0.9943 |

**Table S3. Wilcoxon signed rank test and FDR corrections results.** We computed one-directional Wilcoxon paired tests between pairs of conditions at $alpha = 0.05$: mean RP amplitudes and slopes within each condition for channel FC1, C3, Cz separately and ROI (average across channels ) and mean RP amplitudes obtained through the EMSF technique. The uncorrected p-values were all adjusted with the Benjamini and Hochberg (1995) procedure for controlling the false discovery rate (FDR) for multiple comparison correction. (*) indicates p-values <0.05.

| **Comparisons** | **FC1** | **C3** | **Cz** |
|---|---|---|---|
| 1 vs 2 | (*) | n.s. | n.s. |
| 1 vs 3 | (*) | (*) | (*) |
| 1 vs 4 | (*) | (*) | n.s. |
| 1 vs 5 | n.s. | n.s. | n.s. |
| 2 vs 3 | n.s. | n.s. | n.s. |
| 2 vs 4 | (*) | (*) | n.s. |
| 2 vs 5 | n.s. | n.s. | n.s. |
| 3 vs 4 | n.s. | n.s. | n.s. |
| 3 vs 5 | n.s. | n.s. | n.s. |
| 4 vs 5 | n.s. | n.s. | n.s. |

**Table S4. Cluster-based permutation tests results.** We report the results of the nonparametric statistical tests performed with the method Monte-Carlo on the beta coefficients of the regressions. *cfg.clusteralpha*, *cfg.correcttail*, *cfg.neighbour* correspond to Fieldtrip parameters. *stat.posclusters.prob* is the output of the permutation test for positive clusters. (*) indicates p-values <0.05.(**) indicates p-values <0.005.

| **Regression 1 (RPs)** | **Subjects** | **Latency** | **Positive cluster** | **Negative cluster** |
|---|---|---|---|---|
| All RP subjs vs 0 | $n = 22$ | $-2s\ -0.2s$ | $p = 0.0400\ (*)$ | p= 0.3786 (n.s.) |
| Negative RPs vs 0 | $n = 12$ | $-2s\ -0.2s$ | $p = 0.0300\ (*)$ | $p = 0.4585\ (n.s.)$ |
| Positive RPs vs 0 | $n = 10$ | $-2s\ -0.2s$ | $p = 0.1229\ (n.s.)$ | $p = 0.4835\ (n.s.)$ |
| Timer RP subjs vs 0 | $n = 15$ | $-2s\ -0.2s$ | $p = 0.3027\ (n.s.)$ | $p = 0.0340\ (*)$ |
| Non-timer RP subjs vs 0 | $n = 7$ | $-2s\ -0.2s$ | $p = 0.2148\ (n.s.)$ | $p = 0.6224\ (n.s.)$ |
| All RP subjs vs 0 | $n = 22$ | $-0.2s\ \ 1s$ | $p = 0.3237\ (n.s.)$ | $p = 0.1668\ (n.s.)$ |
| Negative RPs vs 0 | $n = 12$ | $-0.2s\ \ 1s$ | $p = 0.8202\ (n.s.)$ | $p = 0.6094\ (n.s.)$ |
| Positive RPs vs 0 | $n = 10$ | $-0.2s\ \ 1s$ | $p = 0.3027\ (n.s.)$ | $p = 0.0340\ (*)$ |
| Timer RP subjs vs 0 | $n = 15$ | $-0.2s\ \ 1s$ | $p = 0.5874\ (n.s.)$ | $p = 0.5355\ (n.s.)$ |
| Non-timer RP subjs vs 0 | $n = 7$ | $-0.2s\ \ 1s$ | $p = 0.1049\ (n.s.)$ | $p = 9.9900e-04\ (***)$ |
| **Regression 1 (LRPs)** | **Subjects** | **Latency** | **Positive cluster** | **Negative cluster** |
| All LRP subjs vs 0 | $n = 22$ | $-2s\ -0.2s$ | $p = 0.6084\ (n.s.)$ | $p = 0.1349\ (n.s.)$ |
| Negative LRP subjs vs 0 | $n = 12$ | $-2s\ -0.2s$ | $p = 0.5724\ (n.s.)$ | $p = 0.3616\ (n.s.)$ |
| Positive LRP subjs vs 0 | $n = 10$ | $-2s\ -0.2s$ | $p = 0.5305\ (n.s.)$ | $p = 0.7502\ (n.s.)$ |
| Timer LRP subjs vs 0 | $n = 15$ | $-2s\ -0.2s$ | $p = 0.4016\ (n.s.)$ | $p = 0.2557\ (n.s.)$ |
| Non-timer LRP subjs vs 0 | $n = 7$ | $-2s\ -0.2s$ | $p = 0.1968\ (n.s.)$ | $p = 0.5984\ (n.s.)$ |
| All LRP subjs vs 0 | $n = 22$ | $-0.2s\ \ 1s$ | $p = 0.0559\ (n.s.)$ | *no cluster* |
| Negative LRP subjs vs 0 | $n = 12$ | $-0.2s\ \ 1s$ | $p = 0.0130\ (**)$ | *no cluster* |
| Positive LRP subjs vs 0 | $n = 10$ | $-0.2s\ \ 1s$ | *no cluster* | *no cluster* |
| Timer LRP subjs vs 0 | $n = 15$ | $-0.2s\ \ 1s$ | $p = 0.1479\ (n.s.)$ | *no cluster* |
| Non-timer LRP subjs vs 0 | $n = 7$ | $-0.2s\ \ 1s$ | $p = 0.0549\ (n.s.)$ | *no cluster* |
| **Regression 2 (RPs)** | **Subjects** | **Latency** | **Positive cluster** | **Negative cluster** |
| All RP subjs vs 0 | $n = 22$ | $-2s\ -0.2s$ | $p = 0.0270\ (*)$ | $p = 0.4995\ (n.s.)$ |
| Negative RP subjs vs 0 | $n = 12$ | $-2s\ -0.2s$ | $p = 0.0060\ (**)$ | $p = 0.2268\ (n.s.)$ |
| Positive RP subjs vs 0 | $n = 10$ | $-2s\ -0.2s$ | $p = 0.4835\ (n.s.)$ | $p = 0.2507\ (n.s.)$ |
| Timer RP subjs vs 0 | $n = 15$ | $-2s\ -0.2s$ | $p = 0.0030\ (**)$ | $p = 0.3477\ (n.s.)$ |
| Non-timer RP subjs vs 0 | $n = 7$ | $-2s\ -0.2s$ | $p = 0.07899\ (n.s.)$ | $p = 0.5554\ (n.s.)$ |
| All RP subjs vs 0 | $n = 22$ | $-0.2s\ \ 1s$ | $p = 0.2128\ (n.s.)$ | $p = 0.4216\ (n.s.)$ |
| Negative RP subjs vs 0 | $n = 12$ | $-0.2s\ \ 1s$ | $p = 0.3017\ (n.s.)$ | $p = 0.8322\ (n.s.)$ |
| Positive RP subjs vs 0 | $n = 10$ | $-0.2s\ \ 1s$ | $p = 0.3017\ (n.s.)$ | $p = 0.8322\ (n.s.)$ |
| Timer RP subjs vs 0 | $n = 15$ | $-0.2s\ \ 1s$ | $p = 0.6513\ (n.s.)$ | $p = 0.8342\ (n.s.)$ |
| Non-timer RP subjs vs 0 | $n = 7$ | $-0.2s\ \ 1s$ | $p = 0.0639\ (n.s.)$ | $p = 0.2118\ (n.s.)$ |
| **Regression 2 (LRPs)** | **Subjects** | **Latency** | **Positive cluster** | **Negative cluster** |
| All LRP subjs vs 0 | $n = 22$ | $-2s\ -0.2s$ | $p = 0.0110\ (*)$ | $p = 0.3536\ (n.s.)$ |
| Negative LRP subjs vs 0 | $n = 12$ | $-2s\ -0.2s$ | $p = 0.0080\ (*)$ | $p = 0.7952\ (n.s.)$ |
| Positive LRP subjs vs 0 | $n = 10$ | $-2s\ -0.2s$ | $p = 0.2997\ (n.s.)$ | $p = 0.5704\ (n.s.)$ |
| Timer LRP subjs vs 0 | $n = 15$ | $-2s\ -0.2s$ | $p = 0.0480\ (*)$ | $p = 0.4286\ (n.s.)$ |
| Non-timer LRP subjs vs 0 | $n = 7$ | $-2s\ -0.2s$ | $p = 0.4915\ (n.s.)$ | $p = 0.1049\ (n.s.)$ |
| All LRP subjs vs 0 | $n = 22$ | $-0.2s\ \ 1s$ | $p = 0.0060\ (**)$ | *no cluster* |
| Negative LRP subjs vs 0 | $n = 12$ | $-0.2s\ \ 1s$ | $p = 0.0110\ (*)$ | *no cluster* |
| Positive LRP subjs vs 0 | $n = 10$ | $-0.2s\ \ 1s$ | $p = 0.1459\ (n.s.)$ | $p = 0.4496\ (n.s.)$ |
| Timer LRP subjs vs 0 | $n = 15$ | $-0.2s\ \ 1s$ | $p = 0.02209\ (*)$ | *no cluster* |
| Non-timer LRP subjs vs 0 | $n = 7$ | $-0.2s\ \ 1s$ | $p = 0.5225\ (n.s.)$ | $p = 0.1129\ (n.s.)$ |
