## Supplemental Table 2 for "Movement-Preceding Neural Activity under Parametrically Varying Levels of Time Pressure"

**Table S1. Results from one-sample Kolgoromov-Smirnov test.** We tested the normality of the behavioral distribution of the data in each condition. A significant results means that the data is non-normally distributed.

| Condition n° | Statistics | p-values (1.0e-07 *) |
|---|---|---|
| 1 | 0.6014 | 0.0000 |
| 2 | 0.7605 | 0.0000 |
| 3 | 0.8272 | 0.0000 |
| 4 | 0.8744 | 0.0000 |
| 5 | 0.8743 | 0.0000 |

**Table S2. Results from the post-hoc multiple comparison test (Mathworks; Hochberg and Tamhane, 1987).** We run post-hoc pair-wise comparisons comparing the behavioural responses in each condition to all the others conditions in order to reveal where the effect of the time limit conditions were.

| Comparisons | Confidence Interval | | Statistics | p-values |
|---|---|---|---|---|
| 1 vs 2 | − 56.2348 | − 30.0000 | − 3.7652 | 0.0156 |
| 1 vs 3 | − 81.8712 | − 55.6364 | − 29.4015 | 0.0000 |
| 1 vs 4 | − 95.1439 | − 68.9091 | − 42.6743 | 0.0000 |
| 1 vs 5 | − 91.2348 | − 65.0000 | − 38.7652 | 0.0000 |
| 2 vs 3 | − 51.8712 | − 25.6364 | 0.5985 | 0.0592 |
| 2 vs 4 | − 65.1439 | − 38.9091 | − 12.6743 | 0.0005 |
| 2 vs 5 | − 61.2348 | − 35.0000 | − 8.7652 | 0.0025 |
| 3 vs 4 | − 39.5076 | − 13.2727 | 12.9621 | 0.6405 |
| 3 vs 5 | − 35.5985 | − 9.3636 | 16.8712 | 0.8672 |
| 4 vs 5 | − 22.3257 | 3.9091 | 0.1439 | 0.9943 |

**Table S3. Wilcoxon signed rank test and FDR corrections results.** We computed one-directional Wilcoxon paired tests between pairs of conditions at $alpha = 0.05$: mean RP amplitudes and slopes within each condition for channel FC1, C3, Cz separately and ROI (average across channels ) and mean RP amplitudes obtained through the EMSF technique. The uncorrected p-values were all adjusted with the Benjamini and Hochberg (1995) procedure for controlling the false discovery rate (FDR) for multiple comparison correction. (*) indicates p-values <0.05.

| **Comparisons** | **FC1** | **C3** | **Cz** |
|---|---|---|---|
| 1 vs 2 | (*) | n.s. | n.s. |
| 1 vs 3 | (*) | (*) | (*) |
| 1 vs 4 | (*) | (*) | n.s. |
| 1 vs 5 | n.s. | n.s. | n.s. |
| 2 vs 3 | n.s. | n.s. | n.s. |
| 2 vs 4 | (*) | (*) | n.s. |
| 2 vs 5 | n.s. | n.s. | n.s. |
| 3 vs 4 | n.s. | n.s. | n.s. |
| 3 vs 5 | n.s. | n.s. | n.s. |
| 4 vs 5 | n.s. | n.s. | n.s. |

**Table S4. Cluster-based permutation tests results.** We report the results of the nonparametric statistical tests performed with the method Monte-Carlo on the beta coefficients of the regressions. *cfg.clusteralpha*, *cfg.correcttail*, *cfg.neighbour* correspond to Fieldtrip parameters. *stat.posclusters.prob* is the output of the permutation test for positive clusters. (*) indicates p-values <0.05.(**) indicates p-values <0.005.

| **Regression 1 (RPs)** | **Subjects** | **Latency** | **Positive cluster** | **Negative cluster** |
|---|---|---|---|---|
| All RP subjs vs 0 | $n = 22$ | $-2s \quad -0.2s$ | $p = 0.0400$ $(*)$ | p= 0.3786 (n.s.) |
| Negative RPs vs 0 | $n = 12$ | $-2s \quad -0.2s$ | $p = 0.0300$ $(*)$ | $p = 0.4585$ $(n.s.)$ |
| Positive RPs vs 0 | $n = 10$ | $-2s \quad -0.2s$ | $p = 0.1229$ $(n.s.)$ | $p = 0.4835$ $(n.s.)$ |
| Timer RP subjs vs 0 | $n = 15$ | $-2s \quad -0.2s$ | $p = 0.3027$ $(n.s.)$ | $p = 0.0340$ $(*)$ |
| Non-timer RP subjs vs 0 | $n = 7$ | $-2s \quad -0.2s$ | $p = 0.2148$ $(n.s.)$ | $p = 0.6224$ $(n.s.)$ |
| All RP subjs vs 0 | $n = 22$ | $-0.2s \quad 1s$ | $p = 0.3237$ $(n.s.)$ | $p = 0.1668$ $(n.s.)$ |
| Negative RPs vs 0 | $n = 12$ | $-0.2s \quad 1s$ | $p = 0.8202$ $(n.s.)$ | $p = 0.6094$ $(n.s.)$ |
| Positive RPs vs 0 | $n = 10$ | $-0.2s \quad 1s$ | $p = 0.3027$ $(n.s.)$ | $p = 0.0340$ $(*)$ |
| Timer RP subjs vs 0 | $n = 15$ | $-0.2s \quad 1s$ | $p = 0.5874$ $(n.s.)$ | $p = 0.5355$ $(n.s.)$ |
| Non-timer RP subjs vs 0 | $n = 7$ | $-0.2s \quad 1s$ | $p = 0.1049$ $(n.s.)$ | $p = 9.9900e-04$ $(***)$ |
| **Regression 1 (LRPs)** | **Subjects** | **Latency** | **Positive cluster** | **Negative cluster** |
| All LRP subjs vs 0 | $n = 22$ | $-2s \quad -0.2s$ | $p = 0.6084$ $(n.s.)$ | $p = 0.1349$ $(n.s.)$ |
| Negative LRP subjs vs 0 | $n = 12$ | $-2s \quad -0.2s$ | $p = 0.5724$ $(n.s.)$ | $p = 0.3616$ $(n.s.)$ |
| Positive LRP subjs vs 0 | $n = 10$ | $-2s \quad -0.2s$ | $p = 0.5305$ $(n.s.)$ | $p = 0.7502$ $(n.s.)$ |
| Timer LRP subjs vs 0 | $n = 15$ | $-2s \quad -0.2s$ | $p = 0.4016$ $(n.s.)$ | $p = 0.2557$ $(n.s.)$ |
| Non-timer LRP subjs vs 0 | $n = 7$ | $-2s \quad -0.2s$ | $p = 0.1968$ $(n.s.)$ | $p = 0.5984$ $(n.s.)$ |
| All LRP subjs vs 0 | $n = 22$ | $-0.2s \quad 1s$ | $p = 0.0559$ $(n.s.)$ | *no cluster* |
| Negative LRP subjs vs 0 | $n = 12$ | $-0.2s \quad 1s$ | $p = 0.0130$ $(**)$ | *no cluster* |
| Positive LRP subjs vs 0 | $n = 10$ | $-0.2s \quad 1s$ | *no cluster* | *no cluster* |
| Timer LRP subjs vs 0 | $n = 15$ | $-0.2s \quad 1s$ | $p = 0.1479$ $(n.s.)$ | *no cluster* |
| Non-timer LRP subjs vs 0 | $n = 7$ | $-0.2s \quad 1s$ | $p = 0.0549$ $(n.s.)$ | *no cluster* |
| **Regression 2 (RPs)** | **Subjects** | **Latency** | **Positive cluster** | **Negative cluster** |
| All RP subjs vs 0 | $n = 22$ | $-2s \quad -0.2s$ | $p = 0.0270$ $(*)$ | $p = 0.4995$ $(n.s.)$ |
| Negative RP subjs vs 0 | $n = 12$ | $-2s \quad -0.2s$ | $p = 0.0060$ $(**)$ | $p = 0.2268$ $(n.s.)$ |
| Positive RP subjs vs 0 | $n = 10$ | $-2s \quad -0.2s$ | $p = 0.4835$ $(n.s.)$ | $p = 0.2507$ $(n.s.)$ |
| Timer RP subjs vs 0 | $n = 15$ | $-2s \quad -0.2s$ | $p = 0.0030$ $(**)$ | $p = 0.3477$ $(n.s.)$ |
| Non-timer RP subjs vs 0 | $n = 7$ | $-2s \quad -0.2s$ | $p = 0.07899$ $(n.s.)$ | $p = 0.5554$ $(n.s.)$ |
| All RP subjs vs 0 | $n = 22$ | $-0.2s \quad 1s$ | $p = 0.2128$ $(n.s.)$ | $p = 0.4216$ $(n.s.)$ |
| Negative RP subjs vs 0 | $n = 12$ | $-0.2s \quad 1s$ | $p = 0.3017$ $(n.s.)$ | $p = 0.8322$ $(n.s.)$ |
| Positive RP subjs vs 0 | $n = 10$ | $-0.2s \quad 1s$ | $p = 0.3017$ $(n.s.)$ | $p = 0.8322$ $(n.s.)$ |
| Timer RP subjs vs 0 | $n = 15$ | $-0.2s \quad 1s$ | $p = 0.6513$ $(n.s.)$ | $p = 0.8342$ $(n.s.)$ |
| Non-timer RP subjs vs 0 | $n = 7$ | $-0.2s \quad 1s$ | $p = 0.0639$ $(n.s.)$ | $p = 0.2118$ $(n.s.)$ |
| **Regression 2 (LRPs)** | **Subjects** | **Latency** | **Positive cluster** | **Negative cluster** |
| All LRP subjs vs 0 | $n = 22$ | $-2s \quad -0.2s$ | $p = 0.0110$ $(*)$ | $p = 0.3536$ $(n.s.)$ |
| Negative LRP subjs vs 0 | $n = 12$ | $-2s \quad -0.2s$ | $p = 0.0080$ $(*)$ | $p = 0.7952$ $(n.s.)$ |
| Positive LRP subjs vs 0 | $n = 10$ | $-2s \quad -0.2s$ | $p = 0.2997$ $(n.s.)$ | $p = 0.5704$ $(n.s.)$ |
| Timer LRP subjs vs 0 | $n = 15$ | $-2s \quad -0.2s$ | $p = 0.0480$ $(*)$ | $p = 0.4286$ $(n.s.)$ |
| Non-timer LRP subjs vs 0 | $n = 7$ | $-2s \quad -0.2s$ | $p = 0.4915$ $(n.s.)$ | $p = 0.1049$ $(n.s.)$ |
| All LRP subjs vs 0 | $n = 22$ | $-0.2s \quad 1s$ | $p = 0.0060$ $(**)$ | *no cluster* |
| Negative LRP subjs vs 0 | $n = 12$ | $-0.2s \quad 1s$ | $p = 0.0110$ $(*)$ | *no cluster* |
| Positive LRP subjs vs 0 | $n = 10$ | $-0.2s \quad 1s$ | $p = 0.1459$ $(n.s.)$ | $p = 0.4496$ $(n.s.)$ |
| Timer LRP subjs vs 0 | $n = 15$ | $-0.2s \quad 1s$ | $p = 0.02209$ $(*)$ | *no cluster* |
| Non-timer LRP subjs vs 0 | $n = 7$ | $-0.2s \quad 1s$ | $p = 0.5225$ $(n.s.)$ | $p = 0.1129$ $(n.s.)$ |
